# Interferon-gamma induced RNF213 targets nascent double membrane vesicles to inhibit coronavirus primary translation

**DOI:** 10.64898/2026.06.02.729663

**Authors:** Michael A Tartell, Randa Tissawak, Lennard Voss, Manivel Lodha, Christopher Bianco, Masashi Iwamoto, Daniel Poston, Theodora Hatziioannou, Yiska Weisblum, Paul D Bieniasz

## Abstract

Innate immune systems employ many mechanisms to inhibit the replication of viruses that profoundly affect host range and pathogenicity. To identify proteins that restrict human respiratory virus replication, we executed CRISPR screens targeting both type-I and type-II interferon stimulated genes. Among the candidate antiviral genes identified, the giant ring finger protein 213 (RNF213) was a key mediator of antiviral activity of interferon gamma (IFNγ) against the human betacoronavirus HCoV-OC43. RNF213 has ATPase and ubiquitin ligase activities, both of which were required for anti-HCoV-OC43 activity. Using newly designed reporter viruses, we demonstrate that RNF213 acts by a previously undescribed mechanism, inhibiting the translation of incoming viral genomes. RNF213 colocalized and coimmunoprecipitated with the primary viral translation product, NSP3, that orchestrates the formation coronavirus replication compartments. Notably, RNF213 restricted genetically diverse common human coronaviruses, but SARS-CoV-2 apparently evades interaction with and inhibition by RNF213.

---

The recent coronavirus disease 2019 (COVID-19) pandemic, caused by severe acute respiratory syndrome coronavirus 2 (SARS-CoV-2), highlights the importance of understanding respiratory virus pathogenesis and the role of the host proteins in restricting viral replication^1^. The pathogenic impact of coronaviruses varies dramatically; the seasonal human coronaviruses, HCoV-229E, HCoV-NL63, and HCoV-OC43, cause mild upper respiratory tract infections and cold-like symptoms^2^. Conversely, three betacoronaviruses, middle east respiratory syndrome coronavirus (MERS-CoV), SARS-CoV, and most recently SARS-CoV-2^2–5^, are pathogenic in humans and have resulted in local disease outbreaks or pandemics. While the adaptive immune response to coronaviruses has been extensively investigated, the impact of innate immunity is comparatively understudied^3^. The relationship between coronavirus pathogenesis and sensitivity to the innate immune response, and whether this differs between seasonal and pandemic viruses, is unknown. Interferons (IFNs) are a critical component of the innate response to infection and induce the expression of hundreds of interferon-stimulated genes (ISGs)^6,7^. Many ISGs induced by type-I and type-III interferon have been identified, but the role of type-II (IFNγ) specific ISGs in directly restricting viral infection remains poorly described^7–10^.

Following virus entry, the incoming coronavirus (+)RNA viral genome is first translated by host ribosomes as a polyprotein and processed by viral proteases to generate the non-structural proteins (NSPs)^12^. NSP3 (which includes one of the viral proteases) and NSP4 induce the formation of double-membrane vesicles (DMVs), spherules formed by rearrangements of endoplasmic reticulum (ER) derived membranes, which serve as sites of viral RNA synthesis and are a hallmark of positive sense RNA virus replication^13–15^. Though recent studies have increased understanding of DMV structure and formation^17–19^, DMV genesis remain largely enigmatic, and no ISGs that localize to DMVs or interfere with DMV formation are known.

To identify ISGs upregulated by type-I and type-II interferon we performed targeted CRISPR screens in a model epithelial cell line and respiratory viruses from multiple families. Both IFNγ and IFNα potently inhibited the replication of multiple respiratory viruses and numerous ISGs impacted virus replication. Among the ISGs specifically induced by IFNγ, ring finger protein 213 (RNF213) had potent and specific activity against HCoV-OC43. We found that RNF213 inhbits primary translation of the incoming HCoV-OC43 RNA genome and was recruited to nascent virus DMVs. RNF213 inhibited other human coronaviruses HCoV-229E and HCoV-NL63. However, SARS-CoV-2 was resistant to this antiviral mechanism and RNF213 did not associate with SARS-CoV-2 DMVs. Overall, this work reveals an IFNγ-stimulated antiviral gene with a previously undescribed effects on coronavirus translation and DMV formation.

## Results

### IFNγ Induces a larger number of genes than IFNα in a model cell line

To probe the roles of IFNα and IFNγ induced genes in the antiviral response, we generated 20 single-cell clones (SCCs) of A549 cells and measured antiviral activity against a recombinant vesicular stomatitis virus (rVSV) expressing a NanoLuc-luciferase reporter^20^ (Extended Data Fig 1A). Since IFNγ activity varied between clones, we selected one clone (2E10) in which IFNγ exhibited potent anti-VSV activity to probe the sensitivity of four respiratory viruses to IFNα and IFNγ. We tested the (+)RNA human coronavirus HCoV-OC43, the (-)RNA orthopneumovirus respiratory syncytial virus (RSV), the (-)segmented RNA orthomyxovirus influenza A (IAV), and the DNA virus human adenovirus 5 (AdV). While IFNα was more potent than IFNγ, both inhibited infection at sub-nanomolar concentrations (Fig 1A). We performed an RNA-seq experiment in four single-cell clones to identify ISGs upregulated by IFNα and IFNγ mediating antiviral activity. Using 4-fold upregulation as an arbitrary cutoff, IFNγ upregulated a larger number of genes than IFNα (319 vs 136; 119 shared; Fig 1B-C, Extended Data Fig 1B). Overall, 336 genes were upregulated by either IFN type more than 4-fold relative to untreated cells.

**Figure 1:**
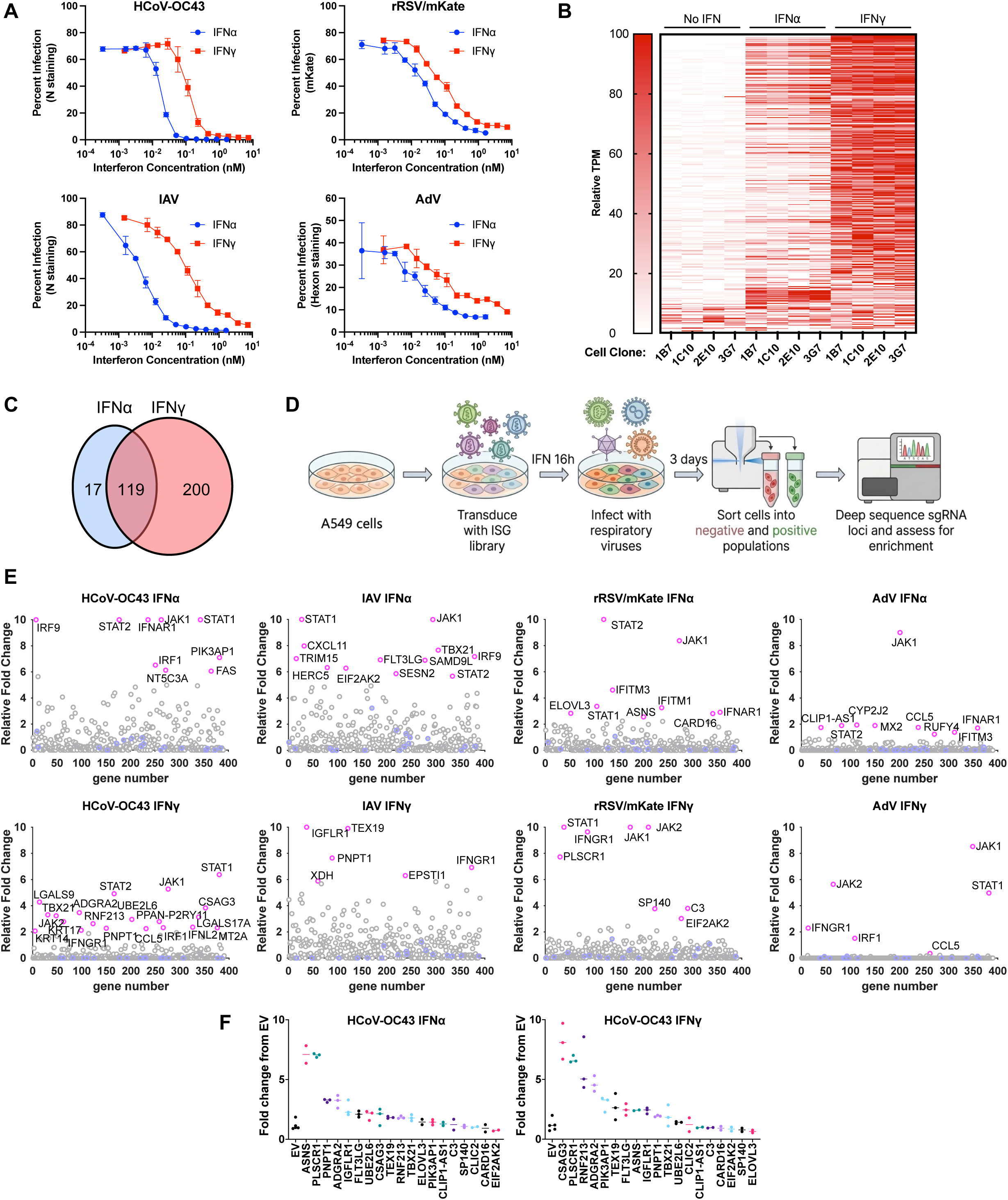
Identification of ISGs that inhibit respiratory virus infection. **(A):** A549 clone 2E10 was treated with IFN⍺/IFNγ for 16h then infected as indicated. Infection was assessed 72 hpi by flow cytometry. **(B):** RNA-seq analysis heatmap depicting normalized expression of selected ISGs across four A549 SCCs (1B7, 1C10, 2E10, 3G7) after 16h mock/IFNα/IFNγ treatment. TPM; transcripts per million. **(C):** Venn diagram of overlap between differentially expressed genes (>4-fold change and padj < 0.05) following IFNα/IFNγ treatment. **(D):** Schematic of screen setup. **(E):** Screen results, where the x axis corresponds to each unique gene in the library, and the y axis denotes the relative fold change. Differentially represented genes in infected versus uninfected populations (p<0.01) are labeled in magenta, non-targeting controls are labeled in blue. **(F):** Flow cytometry of knockout cells treated with IFNα or IFNγ for 16h and infected with HCoV-OC43 for 72h (normalized to EV control).

### IFN-Stimulated Gene Screening Identifies Inhibitors of Respiratory Virus Infection

We used a targeted CRISPR knockout approach to assess the antiviral activity of these 336 ISGs, plus 29 ISGs upregulated following both IFNγ treatment and RSV infection (365 total). We created a targeted single guide RNA (sgRNA) lentiviral library encompassing these ISGs and 100 non-targeting controls and tested them against the aforementioned viruses. Transduced A549 cells (clone 2E10) were incubated with a virus dose that generated ∼1% infection in the context of IFNγ or IFNα inhibition (Extended Data Fig 1C). At 72 hours post infection (hpi), cells were fixed, stained with antibodies to a viral protein, and sorted by flow cytometry into positive and negative populations. Most cells remained negative following IFN treatment and infection, while antigen-positive cells indicated successful infection following sgRNAs targeting of a candidate antiviral ISG (Fig 1D). After sorting, genomic DNA was extracted, sgRNA loci amplified by PCR, and amplicons sequenced to determine sgRNA frequency in infected cells (Extended Data Fig 1D-E). Our screen identified distinct antiviral ISGs for each virus and IFN (Fig 1E, Extended Data Fig 1F). After removing well known hits (e.g. JAK1/STAT1), we selected 19 candidate antiviral genes, generated knockout cells, and infected them with each of the four viruses following IFNα or IFNγ-pretreatment (Fig 1F, Extended Data Fig 1G). Some candidates were recently identified as antiviral (e.g. PLSCR1^21–23^, PNPT1^24^) but others stood out for follow-up, particularly ring finger protein 213 (RNF213). RNF213 is a 591 kDa protein with AAA+ ATPase and E3 ubiquitin ligase domains known for being the major susceptibility gene for Moyamoya disease^25–28^. RNF213 is induced by IFNγ and has extraordinarily broad antimicrobial activity against bacteria, parasites, and DNA viruses^29–34^. As it had never been found to inhibit coronaviruses, we selected its effect on HCoV-OC43 for further investigation.

### RNF213 Inhibits HCoV-OC43 Replication

We knocked out RNF213 in A549 cells using CRISPR-Cas9 and isolated 20 SCCs, as well as 7 empty vector (EV) control SCCs. Cells were pretreated with IFNα or IFNγ for 16 hours, then infected with HCoV-OC43 for 72 hours. RNF213 knockout increased infection by ∼25-fold following IFNγ treatment compared to EV, but had a ≤2-fold effect on HCoV-OC43 infection with no pretreatment or IFNα pretreatment (Fig 2A). Analysis of viral growth-curves showed a similar increase in infectious virus yield upon RNF213 knockout with IFNγ pretreatment, but no effect without IFN (Fig 2B). Quantification of released virions by metabolic labeling revealed an ∼8-fold increase with RNF213 knockout in IFNγ treated cells (Fig 2C, Extended Data Fig 2A), but no changes in the relative proportions of virion proteins (Extended Data Fig 2B). To rule out off-target editing effects, we generated cells with synonymous mutations in the sgRNA target site to disrupt guide recognition. Knock-in of sgRNA-resistance mutations alone did not affect infection, and while RNF213 knockout increased infection with the original sgRNA site, knockout in sgRNA-resistant cells had no effect (Extended Data Fig 2C).

**Figure 2:**
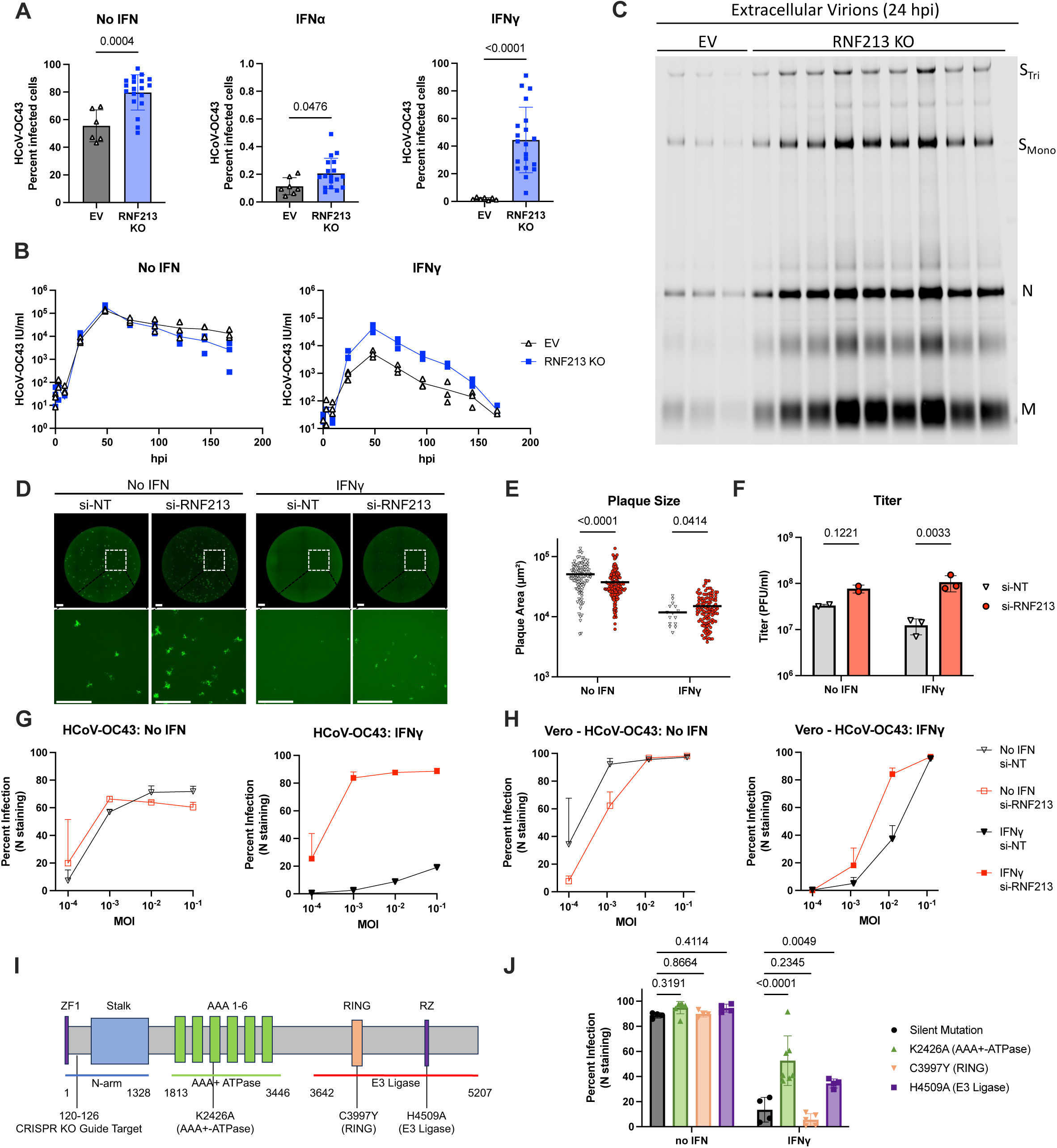
RNF213 restricts HCoV-OC43 replication following IFNγ stimulation. **(A):** RNF213 KO (n=20) or EV (n=7) A549 SCCs were treated as indicated for 16h, infected with HCoV-OC43 at MOI 0.1 for 72h and percent infected cells determined by flow cytometry. Each dot represents an independent SCC (representative experiment, n=3; statistics used each clone as an data point, +/- SD, p values shown: student’s t-test). **(B)**: Growth curve of HCoV-OC43 in A549 EV or RNF213 KO cells pretreated, or not, for 16h with IFNγ and infected at MOI 0.1 (n=3). **(C):** Cells were pretreated with IFNγ for 16h and infected with HCoV-OC43 at MOI 5 for 24h. Labeled extracellular virions were run on an SDS-PAGE gel (representative image, n=3). **(D):** A549 cells were transfected with siRNAs against RNF213 or a non-targeting control (NT), pretreated with IFNγ for 16h, and plaque assay performed for rHCoV-OC43/eGFP (representative image, n=3, scale 1 mm). **(E):** Quantification of plaque size (representative experiment shown +/- SD, p values shown: 2-way ANOVA). **(F):** Quantification of plaque number (n=3, +/- SD, p values shown: 2-way ANOVA). **(G):** siRNA transfected A549 cells were pretreated or not with IFNγ and infected with HCoV-OC43. Percentage infected cells at 72 hpi was determined by flow cytometry (n=3 +/-SD), **(H):** Same as **(G)**, in Vero cells **(I):** Schematic of human RNF213 and locations of CRISPR-mediated editing. **(J):** Flow cytometry of A549 SCCs containing the indicated RNF213 mutations, pretreated with IFNγ (16h) and infected with HCoV-OC43 at MOI 0.1 for 72h. Each dot represents an independent SCC (representative experiment, n=3; statistics based on each clone as an independent experiment, +/- SD, p values shown: 2-way ANOVA). EV: empty vector. KO: knockout. MOI: multiplicity of infection. PFU: plaque forming units. Tri: Trimer. Mono: Monomer. SCC: single-cell clone.

Individual SCCs displayed wide variation in HCoV-OC43 infection (Fig 2A), and some frameshift mutation clones retained low levels of RNF213 protein (Extended Data Fig 2D). To minimize clonal variation, we proceeded using siRNA mediated RNF213 knockdown in unedited cells. A549 cells (clone 2E10) were transfected with siRNA targeting RNF213 or a control siRNA, IFNγ pretreated, and infected with a recombinant HCoV-OC43 expressing an eGFP reporter replacing the nonessential gene NS2^35^ (rHCoV-OC43/eGFP). The siRNA transfection successfully depleted RNF213 (Extended Data Fig 2E) and resulted in an ∼8.6-fold increase in rHCoV-OC43/eGFP titer relative to a control siRNA (Fig 2D-F). IFNγ treatment reduced plaque size, and RNF213 knockdown in this context increased plaque size by ∼1.4-fold (Fig 2E). With wild type HCoV-OC43, RNF213 knockdown caused a 10-fold increase in infection of IFNγ pretreated cells evaluated by flow cytometry at 72 hpi (Fig 2G). We observed a similar (though smaller) effect in Vero and MRC5 cells (Fig 2H, Extended Data Fig 2F). RNF213 knockdown had minimal impact on IAV, RSV, AdV, or VSV infection in any cell type tested (Extended Data Fig 2G). Together, these data demonstrate that in the presence of IFNγ, RNF213 has potent antiviral activity specifically against HCoV-OC43.

To determine which domains and activities of RNF213 mediated its antiviral effect, we knocked-in point mutations to inactivate the RNF213 AAA+ ATPase activity (K2426A), RING domain (C3997Y), or E3 ubiquitin ligase activity (RZ domain, H4509A)^27,32^ (Fig 2I). Knock-in minimally changed protein levels by western blot following IFNγ induction (Extended Data Fig 2H). Mutations that ablated AAA+ ATPase and E3 ubiquitin ligase activities increased HCoV-OC43 infection ∼4-fold and ∼3-fold respectively relative to a synonymous mutation control, while a mutation in the RING domain had no effect (Fig 2J). These data indicate that RNF213 inhbits HCoV-OC43 replication through a mechanism that requires its AAA+ ATPase and E3 ubiquitin ligase activities.

### RNF213 Inhibits HCoV-229E and HCoV-NL63 but not SARS-CoV-2

To determine whether RNF213 restricts other human coronaviruses, we performed infectivity assays with another betacoronavirus, SARS-CoV-2, and the alphacoronaviruses HCoV-229E and HCoV-NL63. An eGFP expressing SARS-CoV-2 reporter virus (rSARS-CoV-2/eGFP) was constructed in which the non-essential ORF7 was replaced with eGFP^36^ (Extended Data Fig 3). HCoV-229E and HCoV-NL63 infection were detected with antibodies against the nucleocapsid protein. For comparison in a matched cellular background, Vero cells or Vero cells expressing human aminopeptidase N (APN, the receptor for HCoV-229E^37^) were used. IFNγ pretreatment had different effects on plaque count and morphology for each virus, causing large reductions in titer but modest reductions in plaque size (e.g. HCoV-229E) or vice-versa (e.g. rSARS-CoV-2/eGFP) (Fig 3A-C). Knockdown of RNF213 in the absence of IFNγ had no effect on any virus tested (Fig 3A-B). RNF213 knockdown with IFNγ pretreatment caused a ∼4-fold increase in titer for rHCoV-OC43/eGFP and HCoV-229E, and a more modest ∼2-fold increase for HCoV-NL63 (Fig 3B). Surprisingly, RNF213 knockdown had a negligible effect on rSARS-CoV-2/eGFP plaque size or titer, despite a ∼5-fold reduction in rSARS-CoV-2/eGFP plaque size with IFNγ pretreatment (Fig 3B-C). We observed similar effects in flow cytometry experiments. RNF213 knockdown increased infection of IFNγ treated cells for rHCoV-OC43/eGFP, HCoV-229E, and to some extent HCoV-NL63, but not rSARS-CoV-2/eGFP (Fig 3D, Extended Data Fig 4). These data show that RNF213 inhibits human coronaviruses HCoV-OC43, HCoV-229E, and HCoV-NL63, but not SARS-CoV-2.

**Figure 3:**
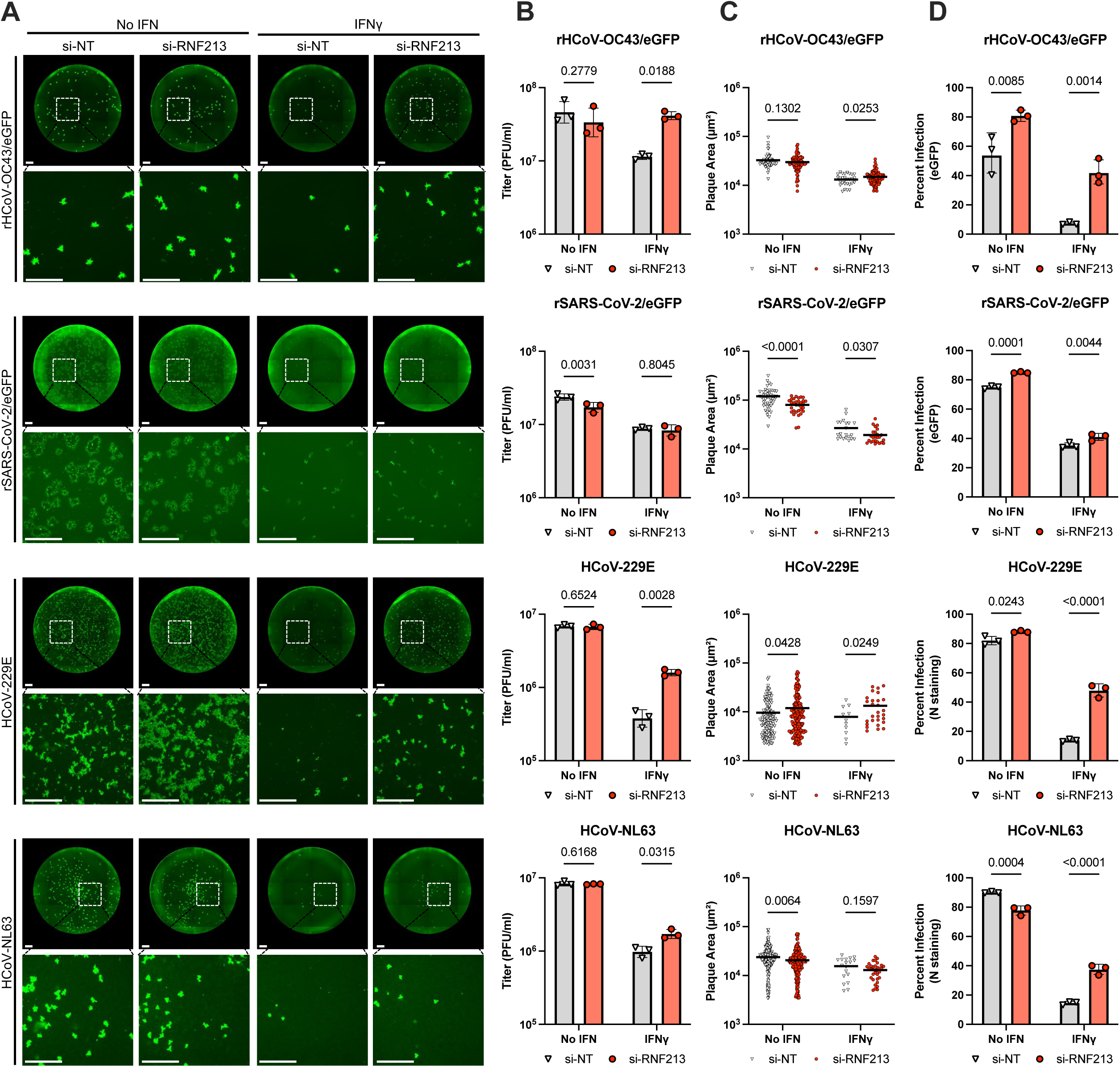
RNF213 inhibits diverse human coronaviruses, but not SARS-CoV-2. Vero cells were transfected with the indicated siRNAs and pretreated with IFNγ for 16 hours. Cells were infected with the indicated virus and subjected to plaque assay (A-C) or flow cytometry (D) analysis. **(A):** Plaque assays, rHCoV-OC43/eGFP and rSARS-CoV-2/eGFP were detected by eGFP reporter expression while HCoV-229E and HCoV-NL63 plaques were stained with antibodies against the corresponding nucleocapsid (representative images, n=3, scale bar: 1 mm). **(B):** Plaques were counted and titers calculated (n=3, +/-SD, p-values shown: 2-way ANOVA). **(C)**: Plaque size was calculated (representative experiment shown, n=3, p-values shown: student’s t-test). **(D):** Flow cytometry was performed at 3 dpi (rHCoV-OC43/eGFP MOI 0.01, HCoV-229E MOI 0.25, HCoV-NL63 MOI 0.8) or 72 hpi (rSARS-CoV-2/eGFP MOI 0.7) (n=3, +/-SD, p-values shown: 2-way ANOVA).

### RNF213 Inhibits Translation of Incoming HCoV-OC43 Genomes

We next determined which step of the viral life cycle was inhibited by RNF213 by examining HCoV-OC43 replication within a single round of infection. First, we determined the impact of RNF213 on expression of reporter encoded by a subgenomic RNA using rHCoV-OC43/eGFP (Fig 4A). Replacement of NS2 did not alter inhibition by RNF213 (Extended Data Fig 5A-B). To restrict rHCoV-OC43/eGFP replication to a single intracellular cycle, we hypothesized normal human serum (NHS) could be used as humans are broadly seropositive for HCoV-OC43^38,39^. When added at 4 hpi, 15% NHS halted rHCoV-OC43/eGFP spread (Extended Data Fig 5C-D). A549 EV or RNF213 knockout SCCs were pretreated with IFNγ, and infected for 24 hours with rHCoV-OC43/eGFP with 15% NHS added at 4 hpi. In both the presence (single-round) or absence (multi-round) of NHS, eGFP positive cells were increased by RNF213 knockout (Fig 4B). Genomic and subgenomic RNAs were similiarly increased (Fig 4C). We thus conclude that RNF213 inhibits replication at or before viral RNA synthesis, rather than late protein synthesis, assembly, or egress.

**Figure 4:**
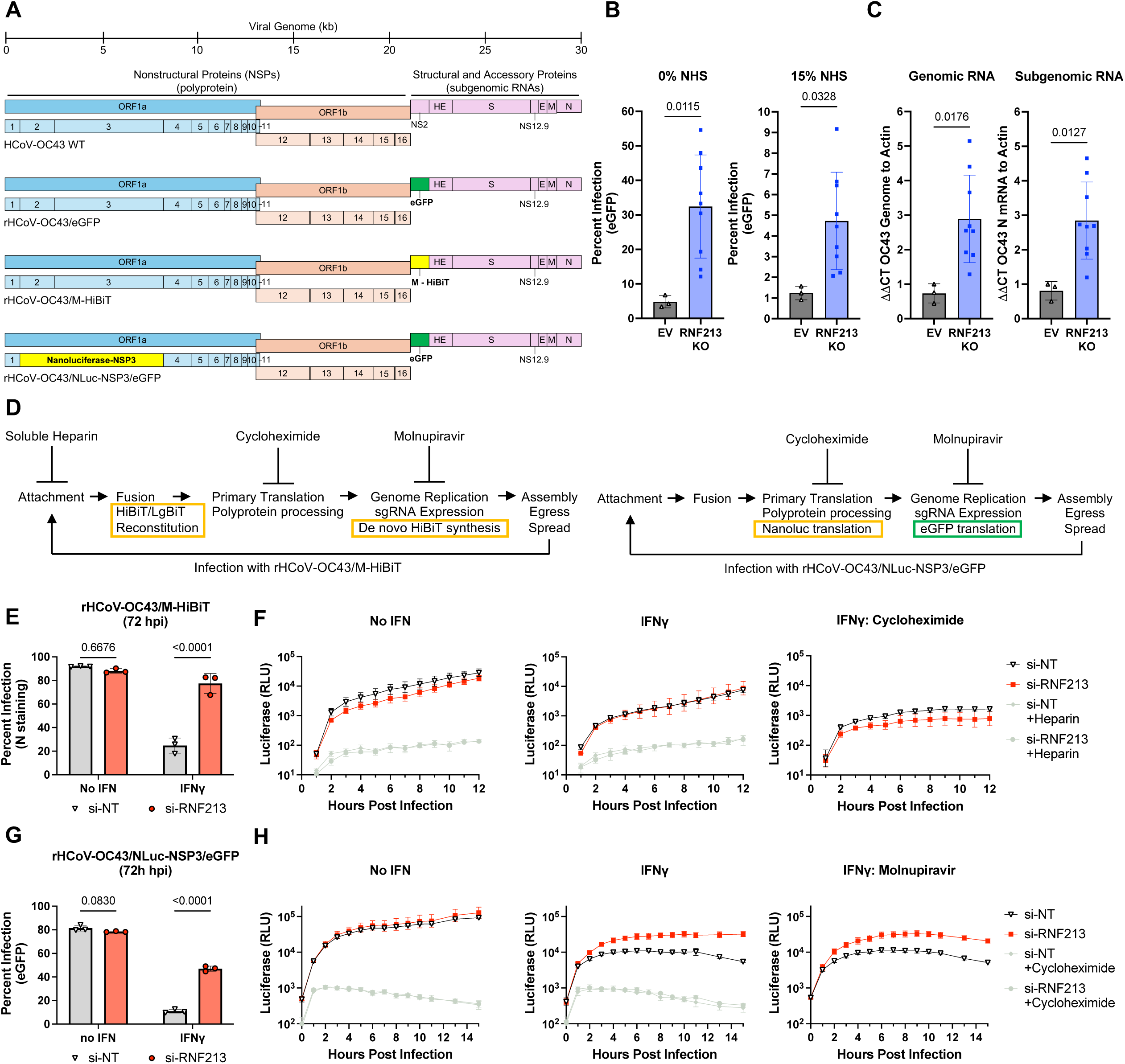
RNF213 inhibits translation of incoming HCoV-OC43 genomic RNA. **(A):** Schematics of recombinant rHCoV-OC43 viruses expressing reporter proteins. **(B):** Flow cytometry analysis at 24 hpi of EV or RNF213 KO A549 SCC cells pretreated with IFNγ for 16h, and infected with rHCoV-OC43/eGFP at MOI 0.1 in the presence or absence of 15% normal human serum added at 4 hpi (NHS) (n=3, +/- SD, p-values displayed, student’s t-test). **(C):** Same as B, except that cells were infected at MOI 1, 15% NHS added at 4 hpi, and qPCR was performed for genomic and subgenomic RNA (n=3, +/- SD, p-values displayed: student’s t-test). **(D):** Schematics of entry (left) and primary translation (right) assays. **(E-H):** Vero cells were transfected with the indicated siRNAs, treated with IFNγ or no IFN for 16 hours, then infected. **(E):** Infection of cells expressing LgBiT with rHCoV-OC43/M-HiBiT at MOI 0.04. Percent infection at 72 hpi determined by flow cytometry (n=3, +/- SD, p-values displayed: 2-way ANOVA). **(F):** Infection of cells expressing LgBiT with rHCoV-OC43/M-HiBiT at MOI 3. Luminescence was measured at the indicated times post infection (n=3, +/- SD). **(G):** Infection with rHCoV-OC43/NLuc-NSP3/eGFP at MOI 0.56. Percent infection at 48 hpi determined by flow cytometry (n=3, +/- SD, p-values displayed: 2-way ANOVA). **(H):** Infection with rHCoV-OC43/Nluc-NSP3/eGFP at MOI 1. Luminescence was detected at the indicated times post infection (n=3, +/-SD).

As viruses with N-terminal NSP3 reporter gene fusions (see below) were attenuated, we performed subsequent experiments in Vero cells that are more permissive to infection. To determine whether RNF213 inhibited viral entry, we constructed a reporter virus containing a second copy of the membrane (M) protein^40^ tagged at the C-terminus with HiBiT, one of two complementing protein fragments (HiBiT/LgBiT) that together reconstitute NanoLuc-luciferase (rHCoV-OC43/M-HiBiT, Fig 4A). Successful fusion of rHCoV-OC43/M-HiBiT virion particles in LgBiT-expressing-cells delivers M-HiBiT to the cytosol, enabling HiBiT/LgBiT complementation and NanoLuc-luciferase activity without new protein synthesis^23^ (Fig 4D). This signal should not be reduced by cycloheximide and should be blocked by heparin, which inhibits HCoV-OC43 attachment (Fig 4D)^41^. Indeed, pre-incubation of HCoV-OC43 with >12.5 mg/mL heparin inhibited infection by >90% (Extended Data Fig 5E), and heparin did not independently block luciferase signal (Extended Data Fig 5F). Molnupiravir, an inhibitor of coronavirus RNA synthesis^42^, was used to ensure reporter activity was not derived from subgenomic RNA synthesis/translation (Fig 5D). Molnupiravir treatment concurrent with infection resulted in a >100-fold reduction in HCoV-OC43 genomic RNA and N subgenomic mRNA as measured by qPCR at 24 hpi and reduced nucleocapsid protein to undetectable levels (Extended Data Fig 5G-H).

**Figure 5:**
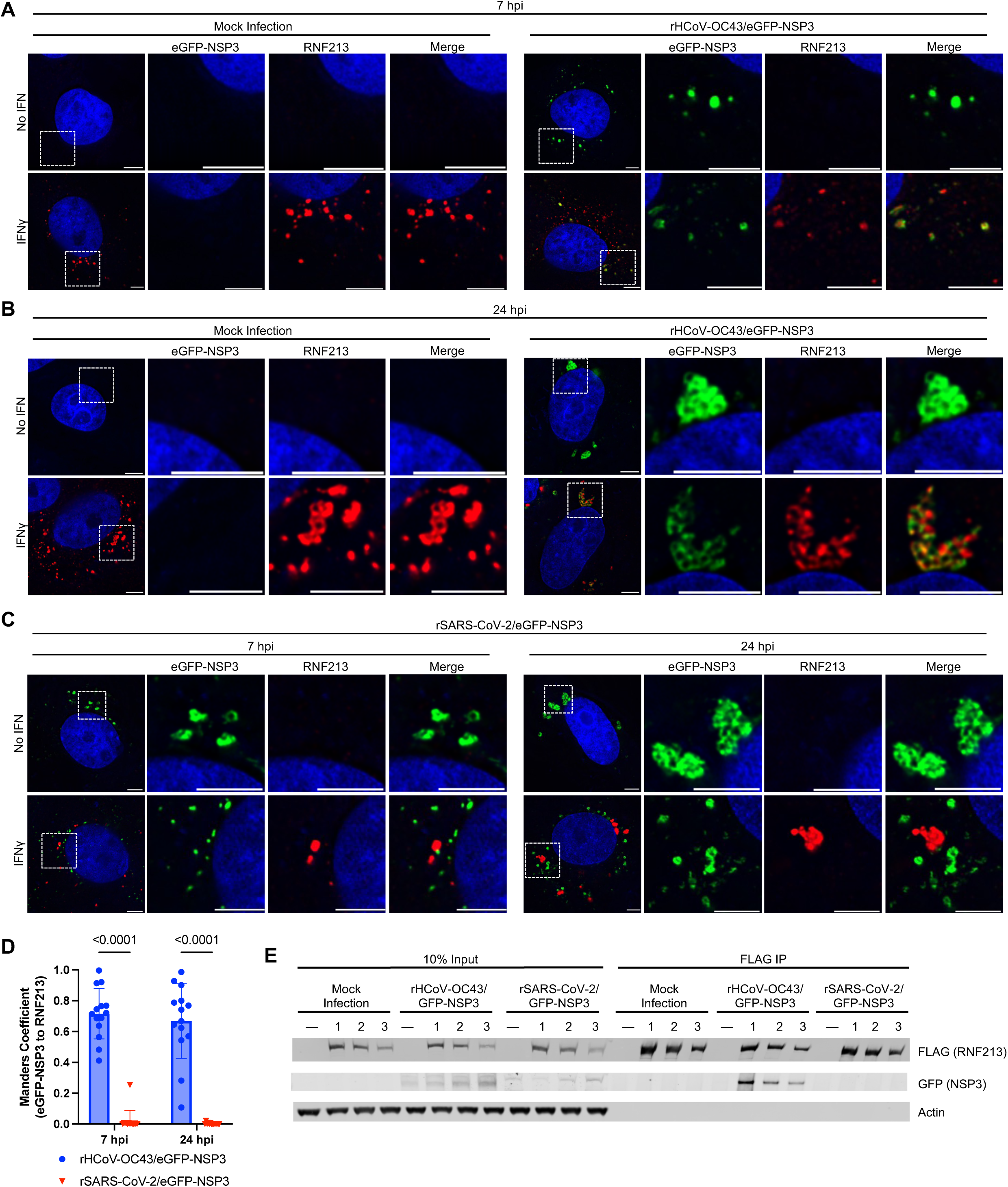
RNF213 recruitment to HcoV-OC43 NSP3, but not SARS-CoV-2 NSP3. Vero cells were pretreated with IFNγ for 24h then mock infected or infected with the indicated virus (green) at MOI 3. Cells were stained for endogenous RNF213 (red) and Hoeschst 33342 (blue). Images shown are a single focal plane from a z-stack (representative images, n=3, scale 5 µm). **(A, B):** Infection with rHCoV-OC43/eGFP-NSP3 at 7 hpi (A) and 24 hpi (B). **(C):** Infection with rSARS-CoV-2/eGFP-NSP3 or rHCoV-OC43/eGFP-NSP3 at the indicated timepoints **(D):** Quantification of the fraction of eGFP-NSP3 signal overlapping with RNF213 (performed for each condition on 14 cells across 3 biological replicates, p-values shown: 2-way ANOVA). **(E):** FLAG-RNF213 Vero cells (3 SCCs) were pretreated with IFNγ for 24h prior to infection at MOI 3 for 24h. FLAG tagged proteins were immunoprecipitated, run on an SDS-PAGE gel, and the indicated protein detected by western blotting (representative images, n=3).

We confirmed that RNF213 inhibited the replication of rHCoV-OC43/M-HiBiT with IFNγ pretreatment at 72 hpi by flow cytometry (Fig 4E, Extended Data Fig 5B) and proceeded to test virus entry. We measured NanoLuc-luciferase activity at one-hour intervals following application of rHCoV-OC43/M-HiBiT virions to LgBiT-expressing target cells. NanoLuc-luciferase activity from HiBiT/LgBiT reconstitution was detected within the first hours of infection at levels >20-fold above the heparin negative control background (Fig 4F, Extended Data Fig 6A). Initial generation of NanoLuc-luciferase activity was not blocked by cycloheximide, indicating that it arose from complementation, not new protein synthesis. We conclude that the impact of RNF213 on HCoV-OC43 is not due to an effect on viral entry. To further determine whether RNF213 affected coronavirus entry, we generated a chimeric, recombinant VSV harboring the spike protein from the RNF213-sensitive coronavirus HCoV-229E, in place of the VSV glycoprotein^43^. rVSV/229E/mNeongreen-P virus enters cells using the entry pathway of HCoV-229E but completes replication using VSV machinery. RNF213 knockdown had no impact on rVSV/229E/mNeongreen-P, nor the parental rVSV/mNeongreen-P (Extended Data 6B), indicating RNF213 does not inhibit entry of rVSV/229E/mNeongreen-P.

To measure translation from the incoming viral genome (primary translation), we constructed a recombinant virus (rHCoV-OC43/NLuc-NSP3/eGFP, Fig 4A) expressing a NanoLuc-luciferase reporter fused to the N-terminus of NSP3 (within the polyprotein), and an eGFP reporter in a subgenomic RNA (Fig 4A, 4D). NSP2 (non-essential for viral replication) was deleted in rHCoV-OC43/NLuc-NSP3/eGFP to avoid increasing viral genome size^44^. RNF213 knockdown increased rHCoV-OC43/NLuc-NSP3/eGFP replication in the presence of IFNγ at 72 hpi, indicating the effect of RNF213 was maintained (Fig 4G, Extended Data Fig 5B). We used cycloheximide treatment concurrent with infection and as a negative control to determine background luciferase levels, and Molnupiravir to block RNA synthesis to ensure that reporter activity was not derived from genome replication. To detect primary translation, we measured NanoLuc-luciferase activity at one hour intervals following infection with rHCoV-OC43/NLuc-NSP3/eGFP (Fig 4H). NanoLuc-luciferase became detectable at 1 hpi, plateauing at 4-7 hpi >50-fold above the cycloheximide control background (Fig 4H). Reporter activity was unaffected by Molnupiravir at these timepoints, indicating its generation did not require viral RNA synthesis and resulted from translation of the incoming genome. IFNγ pretreatment inhibited NanoLuc-Luciferase synthesis ∼4-fold under these conditions and this inhibition was relieved by RNF213 knockdown (Fig 4H). The effect of RNF213 knockdown required pretreatment with IFNγ, as RNF213 knockdown in untreated cells had no effect on NanoLuc-Luciferase synthesis (Fig 4H, Extended Data Fig 6C).

RNF213 accomplishes its antimicrobial effect on bacteria and parasites by ubiquitination and causes degradation of a microbial structure (e.g. LPS ubiquitination and antibacterial autophagy)^29–32^. To determine whether RNF213 altered the stability of NLuc-NSP3, we performed a cycloheximide chase. Control or IFNγ pre-treated cells were infected, then treated with cycloheximide at 7 hpi and the subsequent decay of NLuc-NSP3 was measured (Extended Data Fig 6D-E). RNF213 knockdown did not result in stabilization of NLuc-NSP3 in control or IFNγ pre-treated cells. (Extended Data Fig 6F). We conclude that RNF213 does not induce NLuc-NSP3 degradation. Overall, these experiments indicate that RNF213 restricts coronavirus infection after entry but before the completion of primary translation.

### RNF213 Localizes to HCoV-OC43 DMVs, but Not SARS-CoV-2 DMVs

RNF213 localizes to lipid droplets formed by treatment of cells with oleic acid^27^ and to bacterial or parasitic pathogens during infection^29–32,34^. In uninfected IFNγ-pretreated A549 cells, we found some examples RNF213 co-localized with lipid droplets marked by Lipi-Deep Red, but the majority of RNF213 localized to irregular structures that did not stain with any of the ten markers of cellular organelles tested (Extended Data Fig 7-8). Point mutations in endogenous RNF213 (Fig 2I) changed its localization. Mutating the AAA+ ATPase domain (K2426A) abolished visible structures, while mutating the ubiquitin ligase domain (H4509A) caused RNF213 to localize in smaller puncta even without IFNγ (Extended Data Fig 7C). In IFNγ treated Vero cells, RNF213 localized at both circular and irregular structures (Fig 5A-B), similar to those in A549 cells.

Since RNF213 restricts primary translation of HCoV-OC43 NSP3, and NSP3 is the major structural component of coronavirus DMVs^13,16–19,45^, we examined whether RNF213 is recruited to nascent HCoV-OC43 DMVs. We constructed a virus expressing an eGFP-NSP3 fusion protein (rHCoV-OC43/eGFP-NSP3, Extended Data Fig 3), infected Vero cells for 7 and 24 hpi in the presence or absence of IFNγ, and imaged endogenous RNF213 by immunofluorescence alongside eGFP-NSP3 (Fig 5). Without IFNγ, HCoV-OC43 eGFP-NSP3 localized to small puncta at 7 hpi (Fig 5A), and larger adjoining circular structures at 24 hpi (Fig 5B). In IFNγ-treated rHCoV-OC43/eGFP-NSP3 infected cells, RNF213 co-localized with eGFP-NSP3, at individual puncta at 7 hpi (Fig 5A, Extended Data Fig 9-10) and the larger eGFP-NSP3 structures at 24 hpi (Fig 5B). These data indicate that RNF213 is recruited to HCoV-OC43 DMVs at both early (7 hpi) and late (24 hpi) timepoints.

As SARS-CoV-2 was unaffected by RNF213, we also determined whether RNF213 localized to SARS-CoV-2 DMVs. We constructed a SARS-CoV-2 virus modeled after rHCoV-OC43/eGFP-NSP3 (rSARS-CoV-2/eGFP-NSP3, Extended Data Fig 3) and infected Vero cells. Although both SARS-CoV-2 eGFP-NSP3 and RNF213 were detected in the same cells, they did not co-localize (Fig 5C, Extended Data Fig 9). While ∼70% of the HCoV-OC43 eGFP-NSP3 signal overlapped with RNF213, only ∼2% of SARS-CoV-2 eGFP-NSP3 overlapped with RNF213 (Fig 5D). Similarly, ∼20% (7 hpi) or 40% (24 hpi) of the RNF213 signal co-localized with HCoV-OC43 eGFP-NSP3, but only ∼2% (7 hpi) or <1% (24 hpi) co-localized with rSARS-CoV-2 eGFP-NSP3 (Extended Data Fig 9B).

We next asked if RNF213 could be coprecipitated with either HCoV-OC43 or SARS-CoV-2 NSP3 from infected cells. We knocked-in a N-terminal FLAG tag into the RNF213 locus in Vero cells (3 single cell clones, or null control), which did not affect (FLAG-)RNF213 recruitment to HCoV-OC43 eGFP-NSP3-labelled DMVs (Extended Data Fig 10). IFNγ treated cells were infected with rHCoV-OC43/eGFP-NSP3 or rSARS-CoV-2/eGFP-NSP3, lysates prepared at 24 hpi, subjected to anti-FLAG immunoprecipitation, and both lysates and immunoprecipitates probed for eGFP-NSP3 by western blotting. FLAG-RNF213 co-immunoprecipitated HCoV-OC43 eGFP-NSP3, but not SARS2 eGFP-NSP3 (Fig 5E). Indeed, HCoV-OC43 eGFP-NSP3 was strongly enriched in FLAG-RNF213 immunoprecipitates. Taken together, these data indicate that RNF213 co-localizes and with coprecipitates HCoV-OC43 NSP3, but not SARS-CoV-2 NSP3, concordant with its ability to restrict HCoV-OC43 but not SARS-CoV-2 replication.

## Discussion

Although IFNγ is primarily viewed as an immunomodulatory cytokine^8^, it also has intrinsic antiviral activities. We found that one IFNγ-induced gene, RNF213, is recruited to nascent HCoV-OC43 DMVs and inhibits primary translation of the incoming viral genome. Details of the mechanism by which RNF213 inhibits translation remain to be elucidated. Indeed, facets of the coronavirus primary translation process itself remain poorly understood, as details of how coronaviruses spatially and temporally orchestrate the translation of NSP3 and its protease domain (PLpro), cleavage of NSP1 and NSP2 by PLpro, insertion of NSP3 into the ER, translation and cleavage of NSP4 after insertion, and DMV formation, are unknown^17–19,46–52^. As all these processes may happen concurrently with translation, our findings that RNF213 localizes to HCoV-OC43 DMVs, co-immunoprecipitates NSP3, and inhibits primary translation, suggest a mechanism involving translation arrest on the membranes of nascent DMVs that begin to form as the incoming viral genome is translated on ER membranes.

Diverse coronaviruses (HCoV-OC43, HCoV-229E and HCoV-NL63) are sensitive to RNF213, while SARS-CoV-2 is not. The recruitment of RNF213 to DMVs and the co-immunprecipitation of NSP3 with RNF213 could be the result of direct interaction between RNF213 and NSP3 or other viral proteins. However, prior genetic screens have revealed differences among coronaviruses in host factors required for replication^48–50^ and RNF213 may selectively inhibit certain coronaviruses by sequestering specific required host proteins. The unusual IFNγ-induced RNF213 structures and RNF213 redistribution to DMVs may reflect associations with host factors that mediate DMV formation for some coronaviruses but not others. RNF213 has been reported to target pathogen-associated membrane structures, such as *T. gondii* parasitophorous vacuoles or *S. typhimurium* lipopolysaccaride on the bacterial outer membrane^30–32^, despite the apparent chemical dissimilarities between these membranous compartments. The recruitment of RNF213 to coronavirus DMVs may represent a further example of this phenomenon. The enzymatic activities of RNF213 that are required for its anti-coronavirus activity are also reportedly required for its other antimicrobial effects^25,29,31,32^. However, while RNF213 inhibited these non-viral pathogens by ubiquitination and subsequent degradation of a microbial target^25,30–34^, our results indicate that RNF213 does not cause accelerated degradation of the viral protein NSP3, with which it is associated.

DMVs are a general hallmark of (+)RNA viruses, in addition to coronaviridae^14^. Whether RNF213 can inhibit members of other (+)RNA virus families, or recognizes an element specific to certain members of the coronaviridae, remains to be studied. Indeed, RNF213 may recognize multiple different molecular targets, or target a shared biological pattern or host factor required by certain (+)RNA viruses, bacteria, and parasites.

## Acknowledgments

We thank fellow members of the Laboratory of Retrovirology and the Weisblum Lab for advice and comments. We thank The Rockefeller University Genomics Resource Center for RNA-seq library preparation and sequencing, The Rockefeller University Bioinformatics Resource Center for computational analysis of the RNA-seq datasets, and The Rockefeller University Bio-Imaging Resource Center for computational analysis of plaque size (RRID:SCR_017791). This publication was made possible (in part) with the support of the Charles H. Revson Foundation (to Y.W). The statements made and views expressed, however, are solely the responsibility of the author (Y.W). This work was supported by the Stavros Niarchos Foundation (SNF2025BIE1 to P.D.B), and the Howard Hughes Medical Institute.

## Author Contributions

Conceptualization: M.A.T., Y.W., T.H., P.D.B.; Experiments performed: M.A.T., R.T., L.V., Y.W.; Reverse Genetics Resources: M.A.T., M.L., C.B., M.I.; Screening Resources: D.P., Y.W.; Writing – Original Draft: M.A.T.; Writing – Review & Editing: M.A.T., Y.W., T.H., P.D.B; Supervision: T.H., Y.W., P.D.B.

---

**Extended Data Figure 1:**
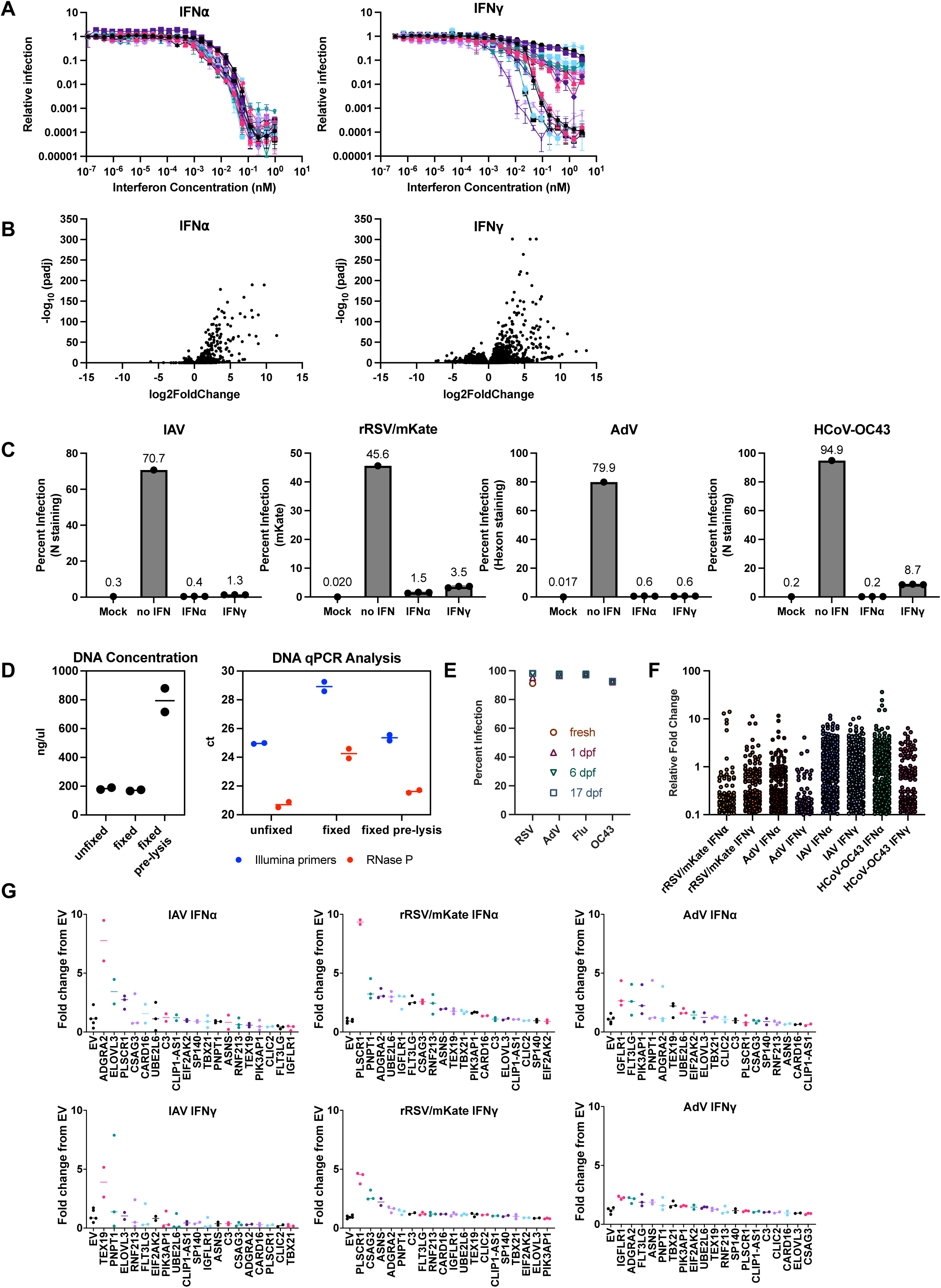
Conditions, validation, and results of CRISPR screen. **(A):** A549 single cell clones (n=20) treated with IFN⍺/IFNγ as indicated for 16h, then infected with rVSV/NanoLuc (MOI 0.003). At 24 hpi cells were lysed and luciferase activity measured. **(B):** RNA-seq analysis of four A549 SCCs (1B7, 1C10, 2E10, 3G7) treated with IFN for 16h. Volcano plots show gene expression changes induced by IFNα or IFNγ compared to mock-treated controls. **(C):** Flow cytometry at 72 hpi of A549 cells (clone 2E10) treated with IFN⍺/IFNγ as indicated for 16 hours, and infected with the indicated virus to produce ∼1% infection after treatment under conditions used for the screen. **(D):** Genomic DNA was extracted from screened cells. DNA concentration was measured, and qPCR performed with the indicated primers to assess the quality of the library. **(E):** Infected A549 cells (clone 2E10) at 3 dpi were fixed for the indicated number of days (dpf: days post fixation), and percent infection was determined by flow cytometry. **(F):** Results of targeted ISG screens against four viruses, analyzed using PinAPL-py. Relative fold change represents the average fold change of guides targeting a gene in the infected population, minus the mean fold change of the control, divided by the standard deviation of the control (sigmaFC). **(G):** Selected ISGs (19) were individually knocked out in A549 cells. Cells were treated with IFNα or IFNγ for 16h then infected with the indicated virus for 72h. Infection levels were measured by flow cytometry and normalized to the empty vector (EV) control.

**Extended Data Figure 2:**
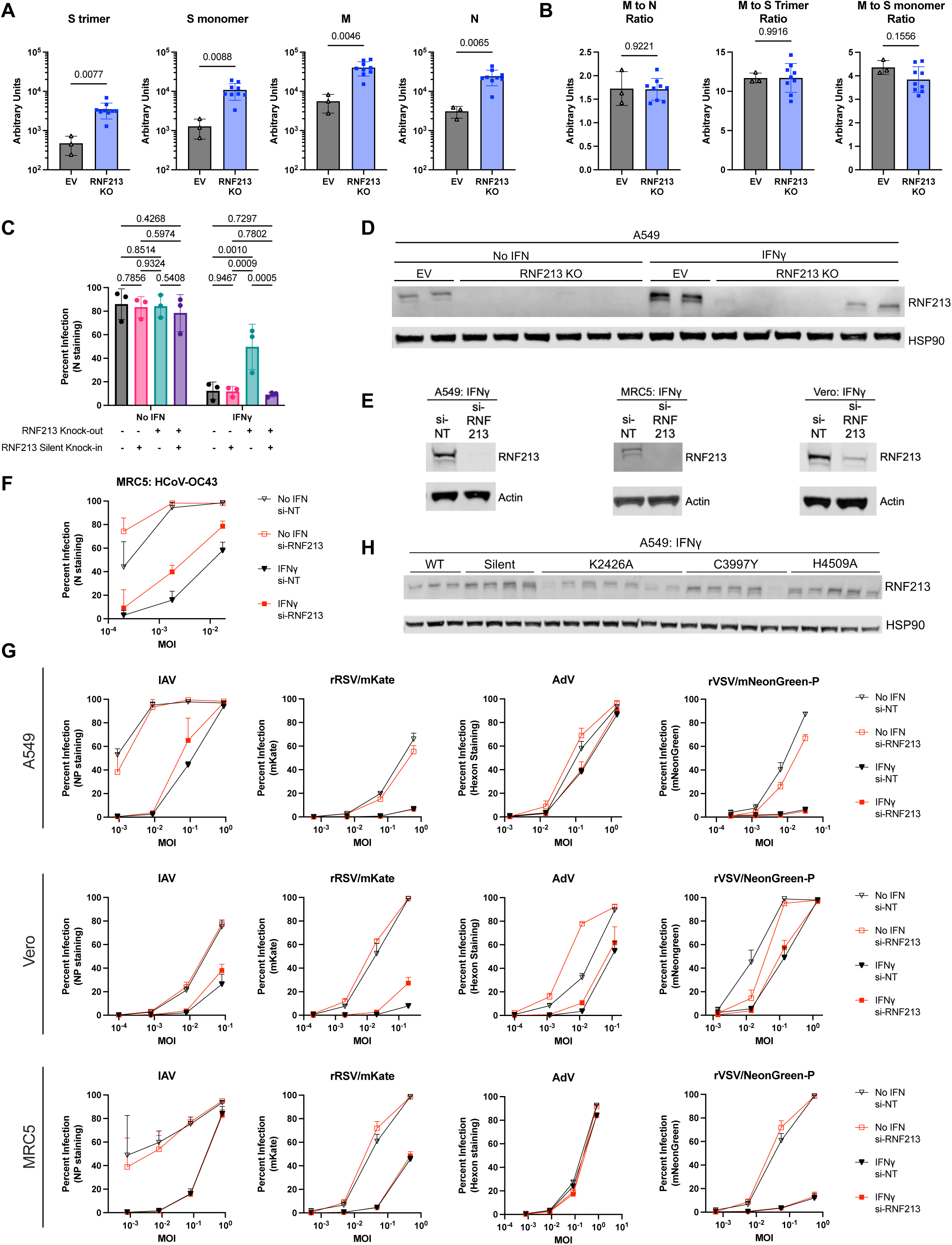
RNF213 restricts HCoV-OC43 replication in multiple cell lines, but not other viruses. **(A):** Quantification of fluorescent signal for each of the indicated bands from Fig 3C (n=3, +/- SD, p-values displayed: student’s t-test). **(B):** Calculation of the ratio of each other protein to M (n=3, +/- SD, p-values displayed: student’s t-test). **(C):** Flow cytometry analysis of A549 RNF213 knock-out cells containing knock-in mutations in the guide RNA target sequence. Cells were pretreated for 16h with IFNγ and infected with HCoV-OC43 at MOI 0.1 for 72 hours (n=3, +/- SD, p-values shown: 2-way ANOVA). **(D):** Western blot of EV or RNF213 KO A549 cells pretreated with IFN for 24 hours (representative images shown, n=3). **(E):** Western blot of the indicated cells transfected with siRNAs against RNF213 or a non-targeting control, then pretreated with IFNγ for an additional 24 hours (representative images shown, n=3). **(F-G):** Flow cytometry of the indicated cells transfected with siRNAs, pretreated with IFNγ for 16 hours, and infected with HCoV-OC43 (F) or the indicated virus (G) at the indicated MOI for 72h, except VSV which was infected for 24h (n=3. +/- SD). **(H):** Western blot of A549 cells with the indicated point mutations pretreated with IFNγ for 24 hours (representative images shown, n=3).

**Extended Data Figure 3:**
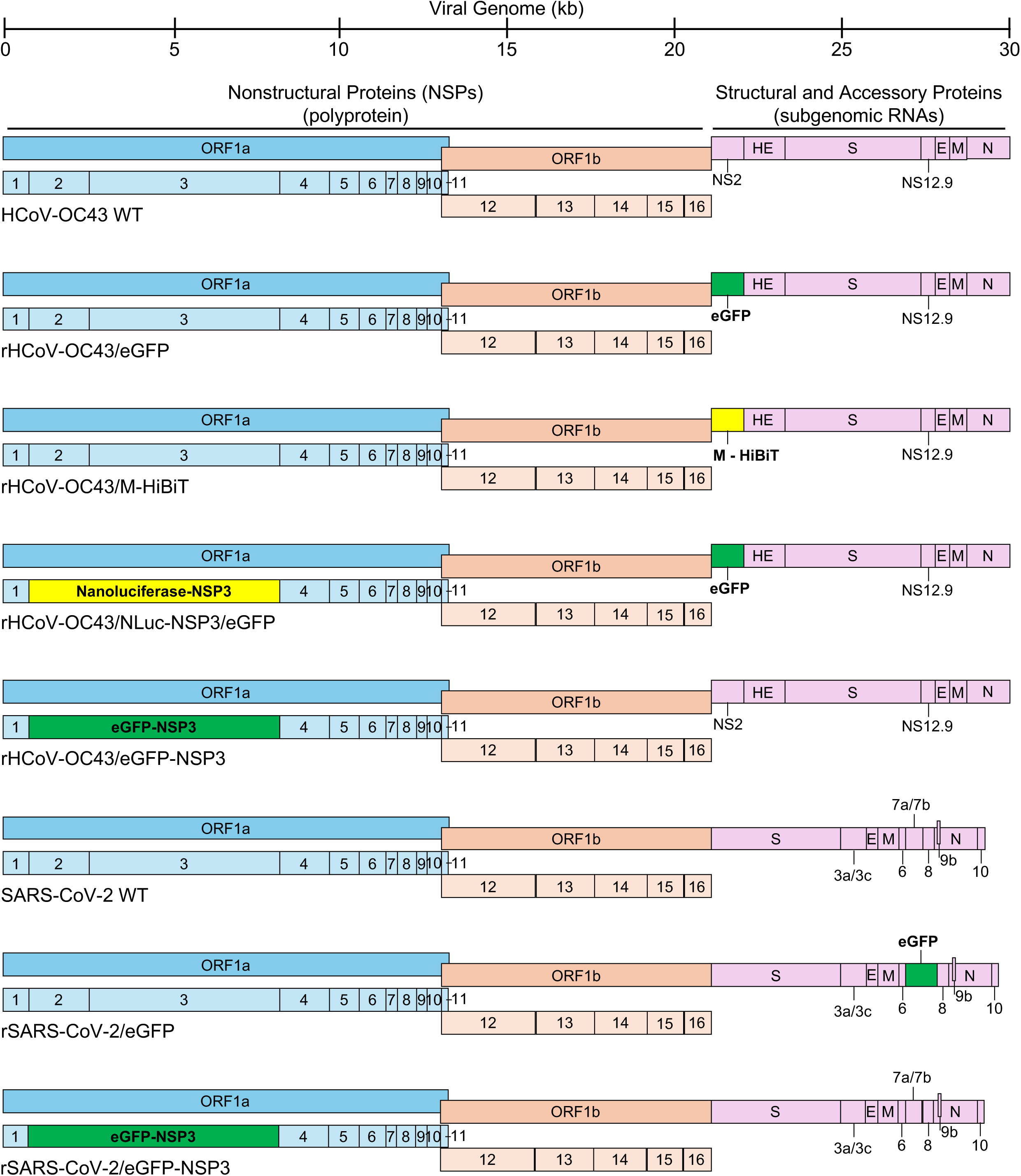
Schematics of recombinant coronaviruses expressing reporter fusions.

**Extended Data Figure 4:**
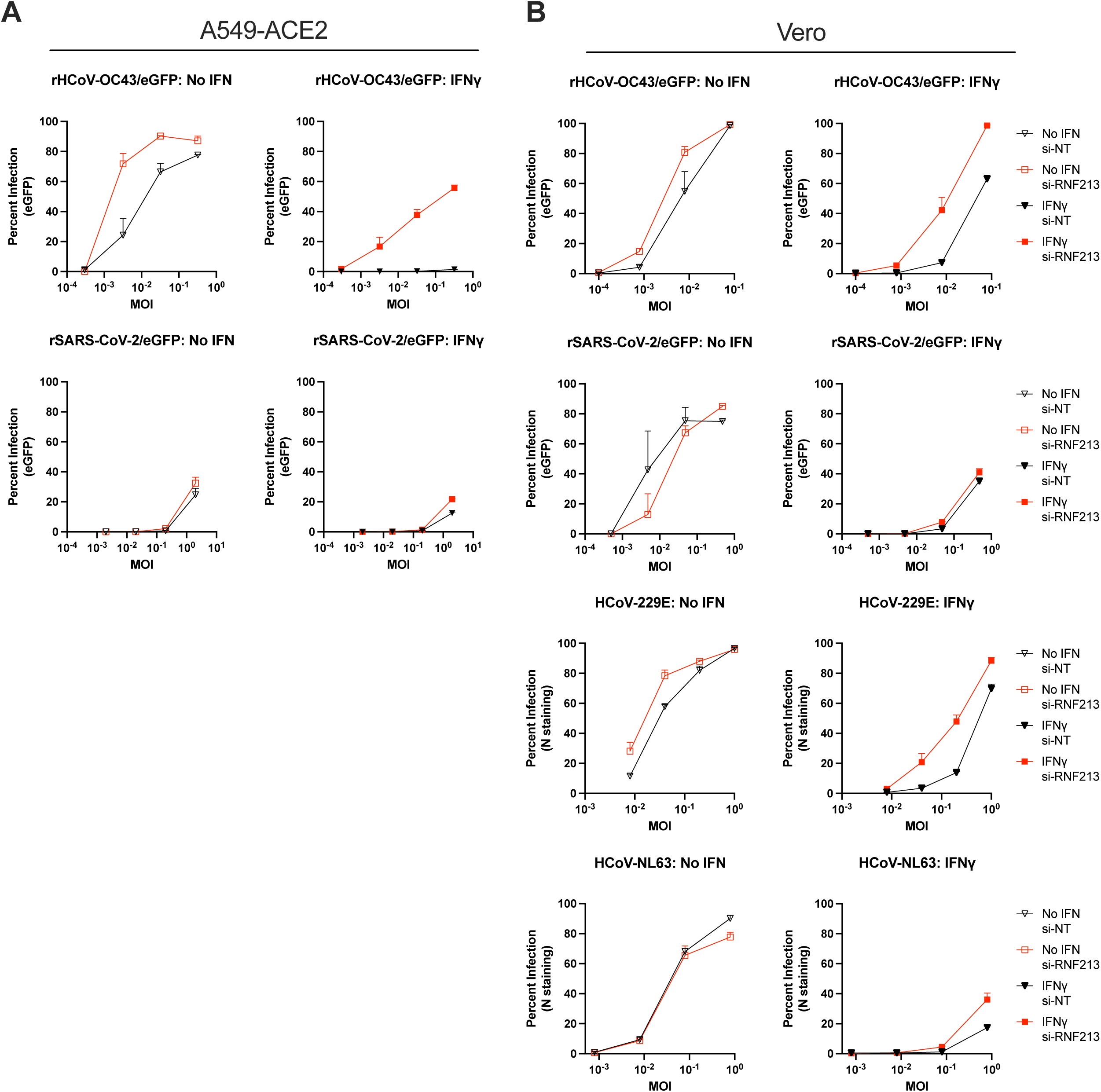
RNF213 restricts diverse human coronaviruses, but not SARS-CoV-2, in multiple cell lines. **(A):** Flow cytometry analysis of A549 cells expressing ACE2 transfected with siRNAs, pretreated with IFNγ for 16 hours, and infected with the indicated virus and MOI for 48h (rSARS-CoV-2/eGFP) or 72h (rHCoV-OC43/eGFP), (n=3. +/- SD). **(B):** Same, in Vero cells. rSARS-CoV-2/eGFP was infected for 48h, all others for 72h.

**Extended Data Figure 5:**
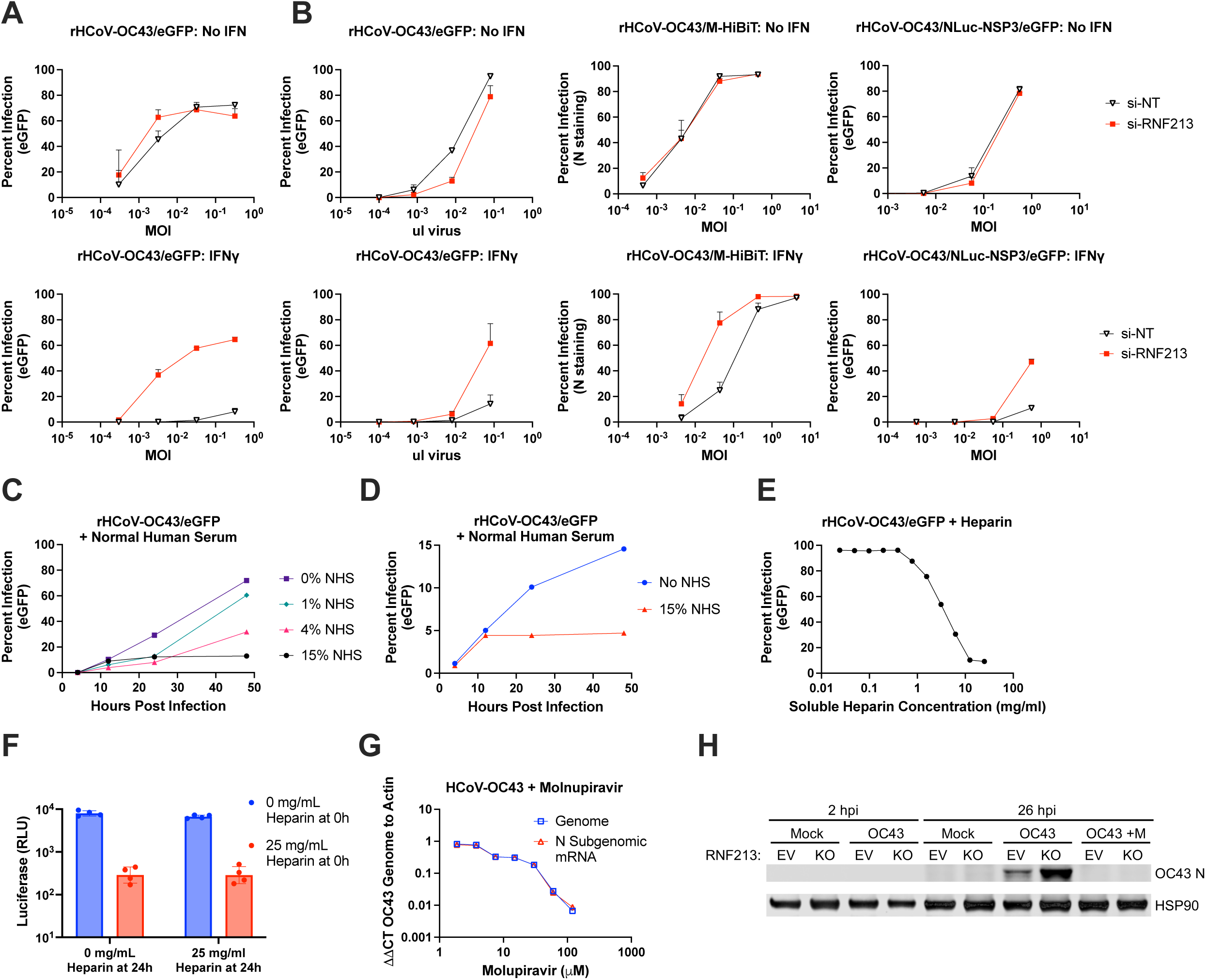
Impact of RNF213 knockdown or inhibitors of steps of the viral life cycle on HCoV-OC43 reporter viruses. **(A):** Flow cytometry analysis of A549 cells transfected with siRNAs, treated with IFNγ for 16 hours, and infected with the indicated virus at the indicated MOI for 72h (n=3, +/- SD). **(B):** Same in Vero cells, except rHCoV-OC43/NLuc-NSP3/eGFP was infected for 48h. **(C):** Flow cytometry analysis of 293T cells infected with rHCoV-OC43/eGFP at MOI 0.1 for 72h, with normal human serum added at 4 hpi (representative experiment, n=3). **(D):** Same with A549 cells. **(E):** Flow cytometry analysis of Vero cells infected with rHCoV-OC43/eGFP at MOI 1 for 24h with soluble heparin added concurrently with infection. **(F):** Vero cells expressing LgBiT were infected with rHCoV-OC43/M-HiBiT at MOI 1 for 24h in the presence of the luciferase detection reagent endurazine. Soluble heparin was added at 0 hpi and/or at 24 hpi, and luminescence detected at 25 hpi (n=3. +/-SD). **(G):** qPCR of A549 cells infected with HCoV-OC43 at MOI 0.1 for 24h in the presence of Molnupiravir (representative images, n=3). **(H):** Western blot of EV or RNF213 KO A549 cells infected for the indicated time with HcoV-OC43 and 200 μM Molnupiravir (representative images, n=3.

**Extended Data Figure 6:**
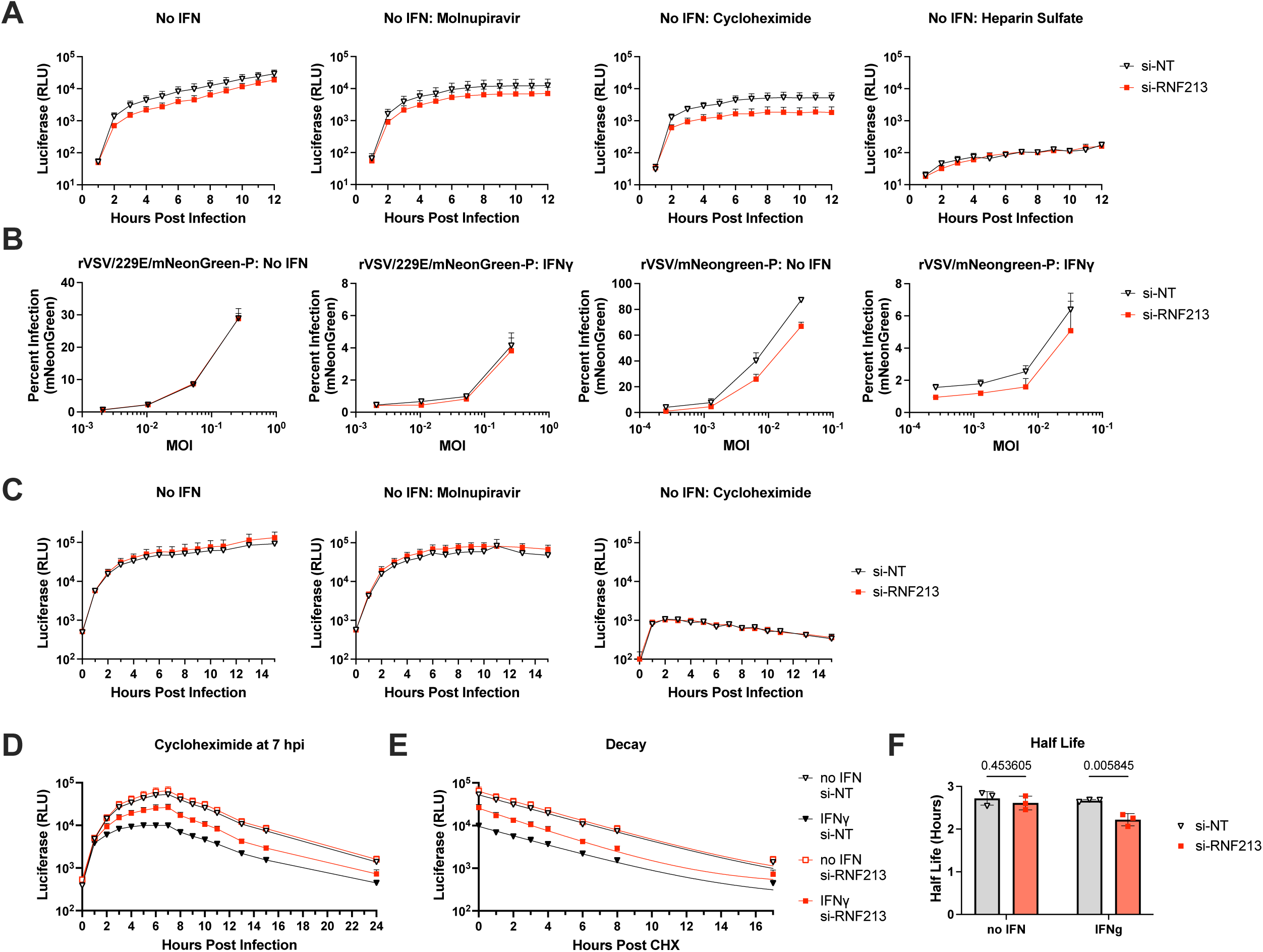
Impact of RNF213 on coronavirus primary translation and entry in the presence or absence of IFNγ. Vero cells were transfected with the indicated siRNAs, treated with IFNγ or no IFN as indicated for 16 hours, then infected with or without the indicated inhibitor concurrently with infection (A, C) or at 7 hpi (D-F) as follows. **(A):** Vero cells expressing LgBiT were infected with rHCoV-OC43/M-HiBiT at MOI 3. Luminescence was detected at the indicated times post infection (n=3, +/- SD). **(B):** Infection with rVSV bearing HCoV-229E spike or the native VSV-G glycoprotein. Percent infection at 24 hpi was determined by flow cytometry (n=3, +/- SD). **(C):** Vero cells were transfected with siRNA, mock treated without interferon for 16 hours, then infected with rHCoV-OC43/NLuc-NSP3/eGFP at MOI 1. Luminescence was detected at the indicated times post infection (n=3, +/- SD). **(D):** Same, except cells were treated with IFNγ for 16h, and cycloheximide was added at 7 hpi. **(E**): The protein decay curve from 7 hpi to 24 hpi from D (equivalent to 0 to 17 hours post cycloheximide treatment) is shown with a nonlinear one phase decay fit line displayed (n=3, +/- SD). **(F):** Calculated half life from the nonlinear fit line from E, n=3, +/-SD, p-value displayed: 2-way ANOVA.

**Extended Data Figure 7:**
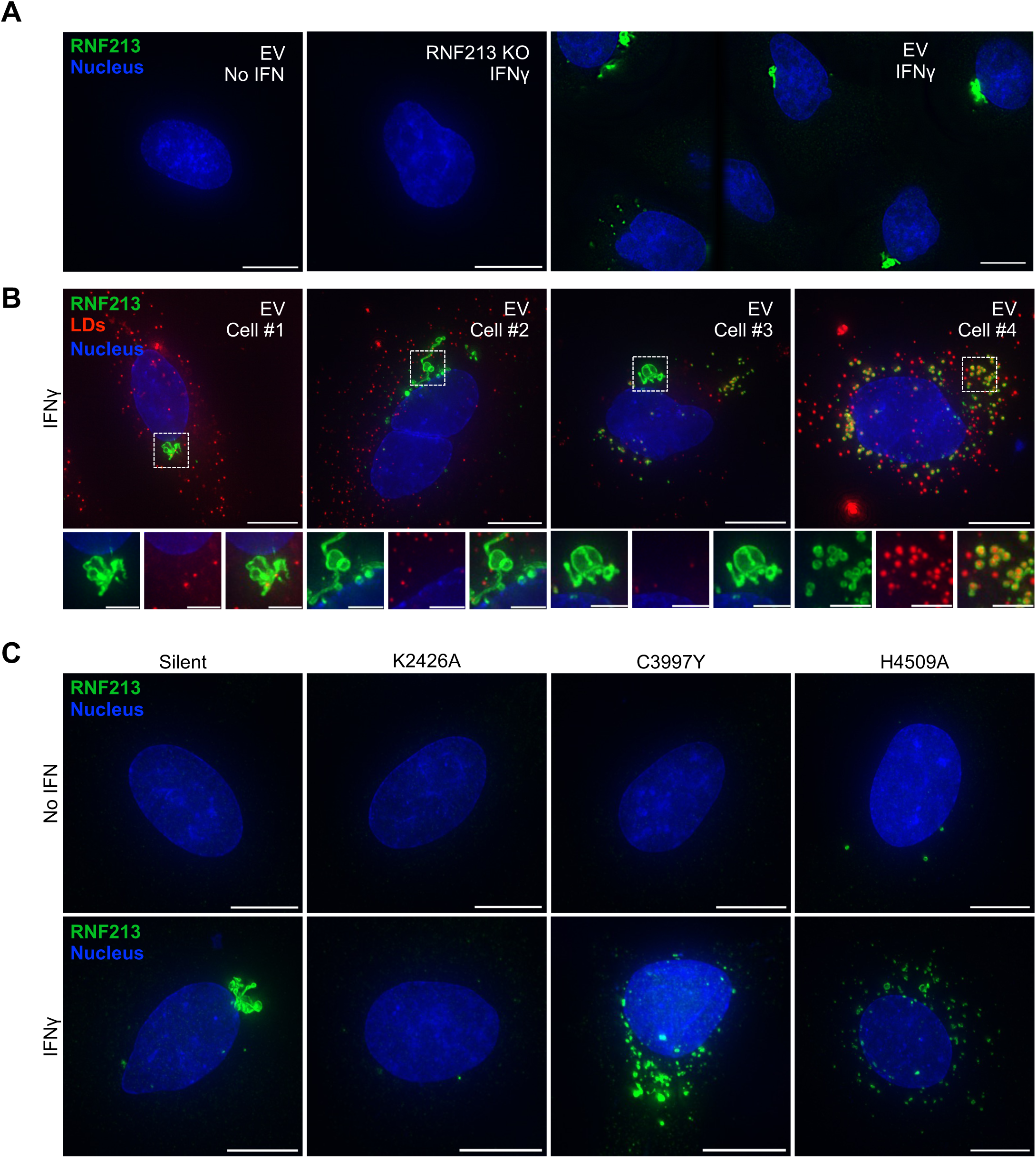
RNF213 localization to irregular subcellular structures following IFNγ stimulation depends on its enzymatic functions. A549 cells were treated with IFNγ for 24 hours and stained for the indicated targets (red, green), and Hoeschst 33342 (blue) for nuclei. **(A):** EV or RNF213 KO cells, stained for endogenous RNF213 (green). Two leftmost images are maximum intensity projection of a z-stack, right image is a selection from a 5x5 mosaic image of a single focal plane. **(B):** Cells cultured with Lipi-Deep Red to stain lipid droplets (red) and stained for endogenous RNF213 (green). **(C):** Cells containing the indicated knock-in mutation in RNF213 stained with an antibody against endogenous RNF213 (green). Except as noted, maximum intensity projection of a z-stack is shown (representative images, n=3; scale bars are 10 µm for large images and 3 µm for magnified insets).

**Extended Data Figure 8:**
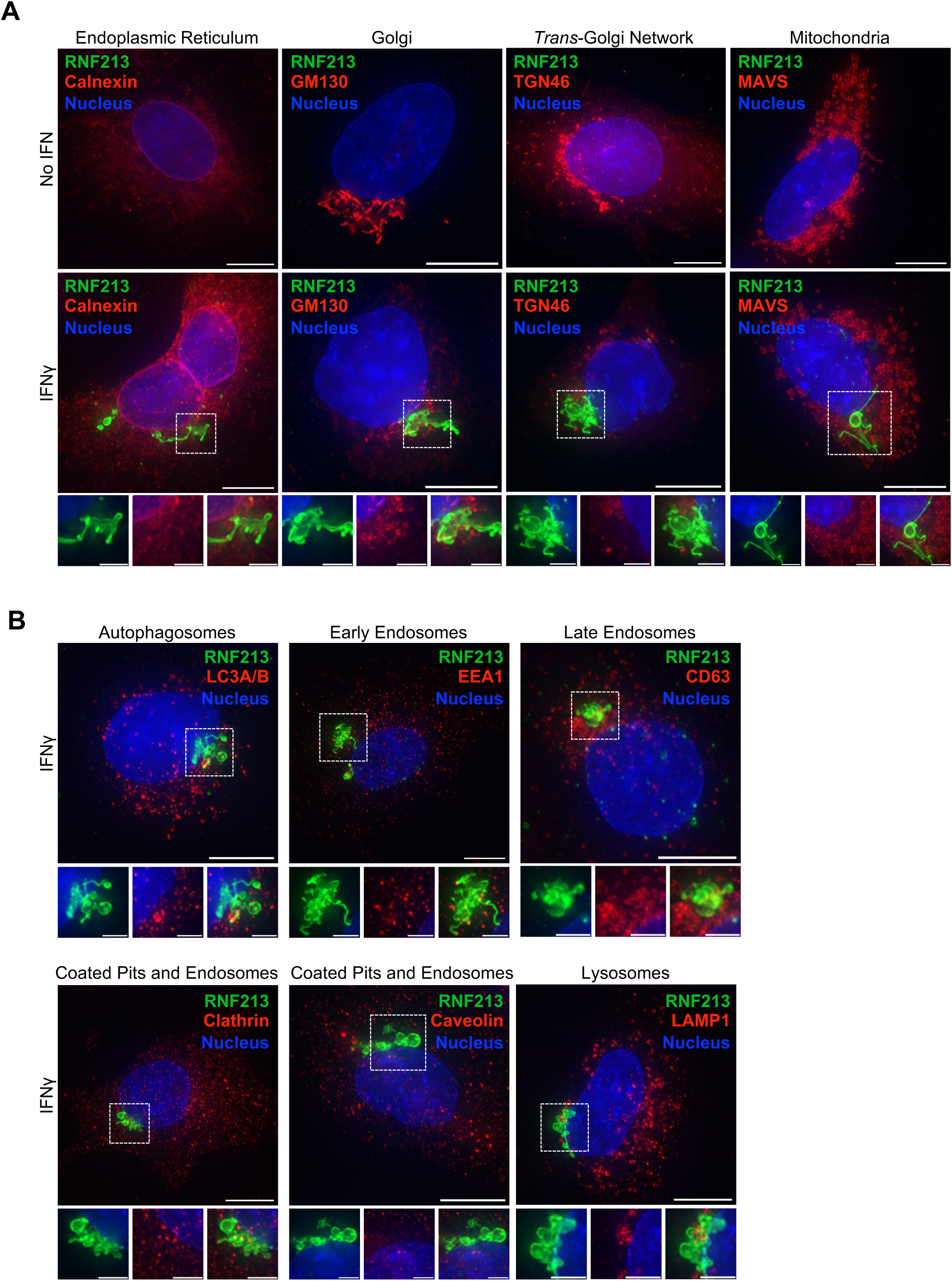
RNF213 does not co-localize with markers of cellular organelles or membrane trafficking following IFNγ pretreatment. EV A549 cells were treated with IFNγ for 24 hours. Cells were stained with Hoeschst 33342 (blue) and antibodies against endogenous RNF213 (green) or the indicated cellular gene (red). **(A):** Cells were stained for known markers of cellular organelles (red). **(B):** Cells were stained for known markers of endosomes or membrane trafficking. Maximum intensity projection of a z-stack is shown, representative images, n=3. Scale bars are 10 µm for large images, and 3 µm for magnified insets.

**Extended Data Figure 9:**
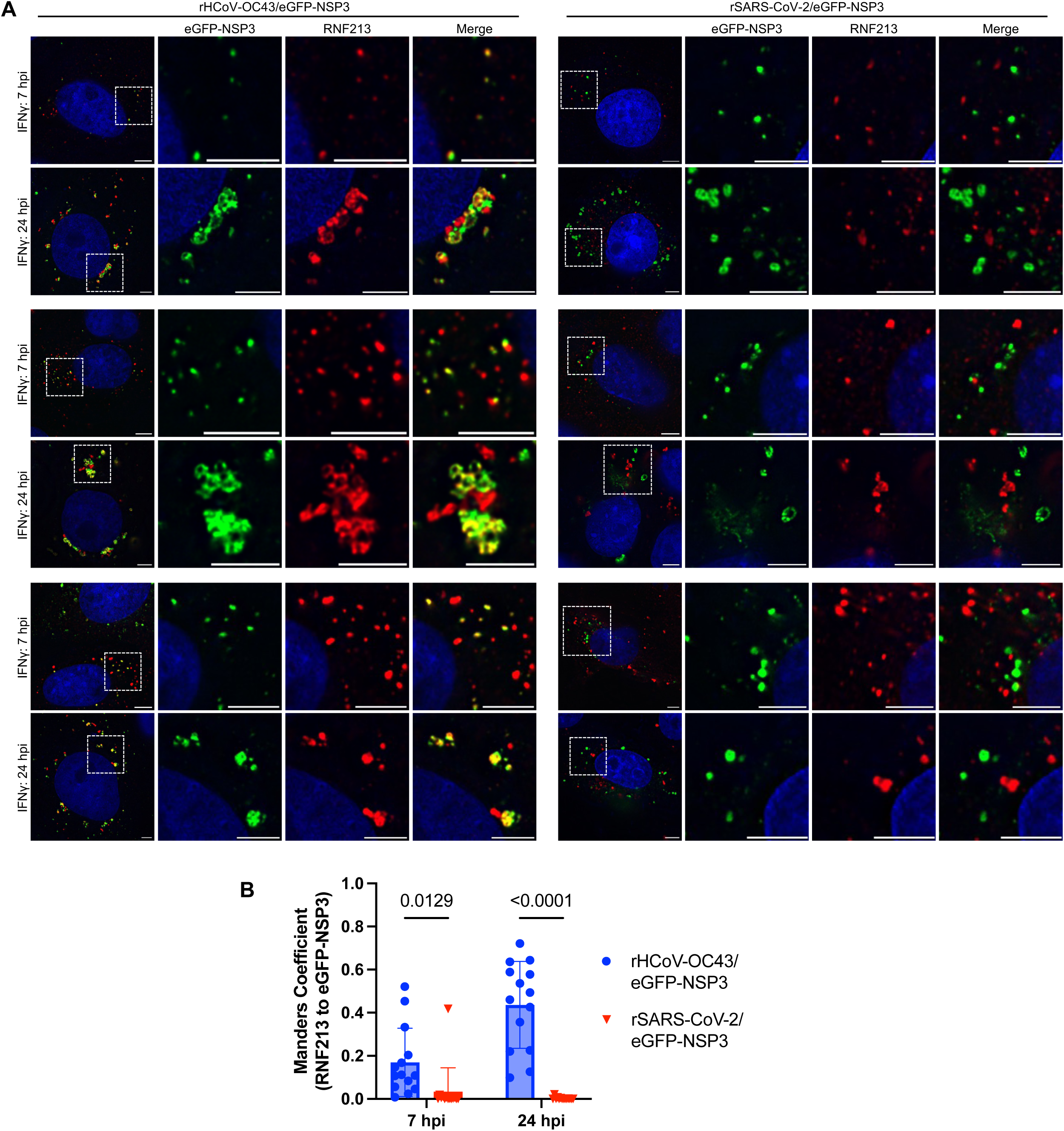
Localization of RNF213 with recombinant coronaviruses expressing reporter fusions. Vero cells were pretreated with IFNγ for 24h, followed by infection with the indicated virus (green) at MOI 3 for the indicated time. Images shown are of a single focal plane in a z-stack, scale bar 5 µm, representative images, n=3. **(A):** Representative images each from 3 separate biological replicates in Vero cells, stained for endogenous RNF213 (red). **(B):** Quantification of the fraction of RNF213 signal overlapping with eGFP-NSP3 (performed for each condition on 14 cells across 3 biological replicates, p-values shown: 2-way ANOVA).

**Extended Data Figure 10:**
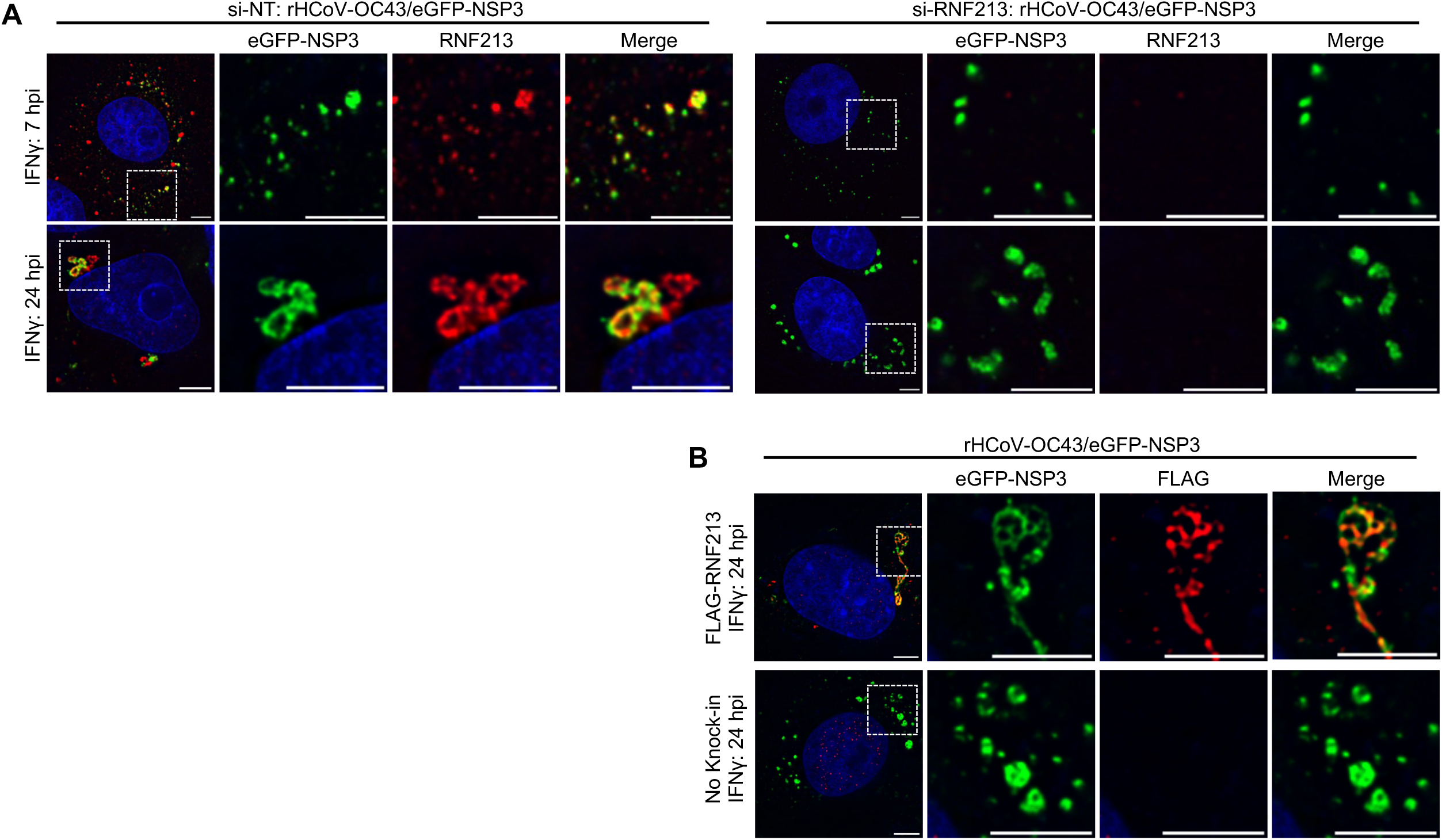
Validation of RNF213 and FLAG antibody staining. Vero cells were pretreated with IFNγ for 24h, followed by infection with the indicated virus (green) at MOI 3 for the indicated time. Images shown are of a single focal plane in a z-stack, scale bar 5 µm, representative images, n=3. **(A):** Vero cells were transfected with siRNAs prior to IFNγ pretreatment and stained for endogenous RNF213 (red). **(B):** Vero cells with an N-terminal FLAG-tag knocked into the endogenous locus of RNF213, or standard Vero cells, stained for FLAG (red).

## Online Methods

### Cell Culture

The cell lines HEK-293T (female human embryonic kidney, ATCC CRL-3216), A549 (male human lung carcinoma, ATCC #CCL-185), MDCK (female *Canis familiaris* dog kidney), MRC5 (male human lung fibroblast, ATCC #CCL-171), Huh7.5 (male human hepatocellular carcinoma, kind gift of Dr. Charles M. Rice), Vero E6 (female *Chlorocebus sabaeus* (African green monkey) kidney, kind gift of Dr. Ralph Baric), and were maintained in Dulbecco’s Modified Eagle’s Medium (DMEM, Thermofisher #11995-065) with 10% fetal calf serum (Sigma #F0926) and 25 μg/mL gentamicin (Thermofisher #15750078) at 37°C and 5% CO_2_. The following antibiotics were used to select transduced cells as needed: puromycin (Sigma #P8833, 1.25 μg/mL), blasticidin (Thermofisher #R21001, 10 μg/mL). Cells were plated for the desired assay in the following formats: 10,000 cells in a 96 well clear plate (Corning #353072) for infectivity and flow cytometry; 25,000 cells in a 96 well black plate for luciferase (Santa Cruz #sc-204450); 20,000 cells in a chamber slide for microscopy (Thermofisher #155409); 100,000 cells in a 24 well plate (Corning #353047) for western blot or plaque assay. Cells were trypsinized by washing in PBS (Thermofisher #21-040-CV) followed by incubation for 5 minutes at 37°C with 0.05% trypsin (Thermofisher #25300-054).

### Production of Sucrose-Cushion Purified Virus Stocks

SARS-CoV-2 viruses (original reference sequence NC_045512.2) were propagated in Vero cells. HCoV-OC43 (ATCC VR-1588) and HCoV-229E (ATCC VR-740) were obtained from Zeptometrix Corporation. HCoV-OC43 was propagated in 293T cells, and HCoV-229E in Huh-7.5 cells. HCoV-NL63 (strain, Amsterdam I) was obtained from the Biodefense and Emerging Infections Research Resources Repository and propagated in Vero cells. Adenovirus 5 was purchased from ATCC (VR-1516) and propagated in A549 cells. Influenza A virus strain A/WSN/33 (H1N1) were propagated in MDCK cells. RSV strain A2-line19F expressing the red fluorescent protein monomeric Katushka 2 (mKate2)^1^ was propagated in Vero cells. VSV viruses (Indiana strain) were propagated on 293T cells. Each virus stock was titered by plaque assay on each cell line used in infectivity experiments separately (A549, Vero, MRC5). Coronavirus stock generation and infections were performed at 34°C, all other viruses were performed at 37°C. When each virus produced visible cytopathic effect in infected cells, viral supernatants were harvested, initially clarified by centrifugation at 1000 x g for 5 minutes, and supernatant further clarified by 0.22 μm sterile filtration (Sigma #S2GPU05RE). Virions were then purified by ultracentrifugation on a 30% sucrose cushion. In a 38.5 ml ultraclear tube (Beckman Coulter # 344058), 9 mL of 30% sucrose was layered under 29.5 mL of viral supernatant, and centrifugated in a SW-32 Ti (Beckman Coulter #369650) swinging-bucket rotor at 25,000 RPM for 1.5 hours at 4 °C. Supernatant was aspirated, and viral pellet resuspended in 3 ml serum-free DMEM.

### Lentivirus Generation and Transduction

Lentivirus proviral particles were generated using an HIV-1_NL4-3_ derived system. 293T cells were transfected with 1 μg of a plasmid encoding HIV NL4-gag-pol, 200 ng of plasmid encoding VSV-G, and 1 μg of plasmid containing a sequence to be integrated and expressed (e.g. pLenti-CRISPRv2) using PEI. The media was changed at 24 hours post transfection (hpt), and supernatants harvested at 48 hpt, filtered through a 0.22 μm syringe filter (Millipore #SLGVR33RS), and used to transduce target cells of interest. At 48 hours post transduction, the media was changed and a selection marker added (e.g. puromycin or blasticidin), and cells selected for a minimum of 3 days.

### RNAseq Experiment and Analysis

A549 cell clones 1B7, 1C10, 2E10, and 3G7 were treated with 1000 U IFNα or IFNγ for 16 h. Total RNA was isolated using the NucleoSpin RNA kit (Macherey-Nagel). RNA-seq libraries were generated from poly(A)-enriched RNA and sequenced at the Rockefeller University Genomics Resource Center on an Illumina NextSeq platform using the High Output 75-cycle kit and standard Illumina sequencing primers. Sequencing data were analyzed by the Rockefeller University Bioinformatics Resource Center, and differential gene expression was determined using DESeq2. Quality control included evaluation of mapping and read assignment metrics.

### ISG library preparation and screening

The library comprised 374 genes represented by 6 sgRNAs per gene, together with 100 non-targeting control sgRNAs. Guides were selected from the Brunello^2,3^ and human GeCKOv2 libraries^4,5^. For genes represented by fewer than 6 unique sgRNAs, additional guides were selected using the Broad GPP sgRNA design tool to ensure uniform representation. The pooled oligonucleotide library (Twist Bioscience) was cloned into the Cas9-expressing lentiviral vector LCRV2. Library representation was confirmed by next-generation sequencing, showing 93.4% perfect-match guides, no undetected guides, and a skew ratio of 4.61.

A549 cells (clone 2E10) were transduced with lentiviral particles carrying the sgRNA library at an MOI of 0.3 and 500× coverage to favor single-guide integration. Cells were treated with IFN alpha or gamma for 16 h and infected with four respiratory viruses. At 3 days post-infection, infected and uninfected/resistant cell populations were isolated by fluorescence-activated cell sorting based on reporter expression (RSV) or antibody staining (AdV, IAV, and HCoV-OC43).

Genomic DNA was extracted using the DNeasy Blood & Tissue Kit (Qiagen) according to the manufacturer’s instructions, with the addition of an overnight proteinase K pre-lysis step at 56°C. Sequencing libraries were prepared by PCR amplification of sgRNA loci as described by Joung et al^6^. Libraries were gel-purified, pooled, and sequenced sequenced on an Illumina NextSeq platform using the High Output kit with 80 cycles for Read 1 and 8 cycles for Index 1 with standard Illumina sequencing primers. Sequencing data were analyzed with PinAPL-py^7^ to identify enriched or depleted sgRNAs and genes relative to the corresponding control populations.

### Generation of Cell Lines by CRISPR Editing

#### Knockout of RNF213

pLentiCRISPRV2 (Addgene plasmid #52961) was used for CRISPR knockout. The guide RNA sequence TGACTTTGCTTTCAAACCCG was cloned by BsmB1 digest of the vector according to the included protocol (https://www.addgene.org/52961/#:∼:text=lentiCRISPRv2%20and%20lentiGuide%20oligo%20cloning%20protocol). Lentivirus particles were produced as above, and transduced cells selected with puromycin before single-cell cloning.

#### Knock-in of Mutations or Tag

Mutations or tag insertion into RNF213 were done using a homology-directed repair method as described by Integrated DNA Technologies (IDT) in which purified Cas9, a guide RNA, and a donor DNA containing the target sequence were nucleofected into cells using a Lonza 4D-Nucleofector X Unit (Lonza # AAF-1003X). A small CRISPR RNA (crRNA) was purchased from IDT for the target locus in RNF213 as follows: Silent mutations in the knockout site: TGACTTTGCTTTCAAACCCG; K2426A: AAACTGGCTGTGGGAAAACC; C3997Y: TCTCCCAGGCAGATGGAGCA; H4509A: CACGGAGCAAGGATGGCCGT; N-terminus: TGGCACGAAGGACACTCCAT. When annealed to a tracrRNA (IDT # 1072533), they form a fully functional sgRNA. A donor DNA was also purchased containing the mutation of interest and at least 35 bp of homology arms (in addition to the gRNA sequence) on either side. First, the crRNA and tracrRNA were mixed by adding 2 μL of a 100 μM stock each and annealed in a thermocycler by incubating at 95 °C for 5 minutes, followed by ramping down to 20 °C at a rate of 0.1 °C per second. 150 pmol of this gRNA mix were added to 125 pmol of Cas9 enzyme and incubated for 20 minutes. Cells were nucleofected using an SF Cell Line 4D-Nucleofector kit (Lonza #V4XC-2024). Cells were trypsinized, spun down at 100 x g for 10 minutes, and 500,000 cells resuspended in 93 μL reconstituted nucleofector solution from the kit. To these cells were added the Cas9 and sgRNA RNP complex, 1.2 μL of 100 μM DNA donor oligo, and 1.2 μL Cas9 enhancer oligo (IDT #1075916). This mixture was nucleofected according to the manufacturer’s preset program for each cell line (A549 or Vero), then incubated for 10 minutes. This mixture was then replated in 10% FCS DMEM containing 1.7 μL/mL HDR enhancer (IDT #10007921), and editing proceeded for 3 days.

#### Selection of Single-Cell Clones

After editing, single-cell clones were isolated by replating cells at a concentration of 0.75 cells / 200 μL, and 200 μL plated in each well of a 96 well plate. After at least a week, cells were visually inspected for single colonies, then trypsinized and expanded.

#### Verification of Editing

Genomic DNA was extracted from cells using QuickExtract Solution (Biosearch Technologies #QE09050) according to manufacturer’s instructions, then diluted 1:1 with water. 5 μL was used to amplify a genomic region of ∼400 bp around the targeted locus for gene editing. The PCR product from each single-cell clone was then re-amplified through PCR so that each individual clone contained a unique barcode. These barcoded PCR products from individual clones were then pooled, gel extracted, subjected to sequencing using an Illumina MiSeq Nano, and analyzed for successful CRISPR editing using Geneious.

### siRNA Transfection

siRNAs were reverse transfected into cells (A549s, Veros, or MRC5s) in either a 12-well format (300,000 cells) or 6-well format (600,000 cells). For a 12-well well, 12.5 pmol siRNA were added to 50 μL Opti-MEM (Thermofisher #31985062) in one tube, and 4 μL RNAiMAX transfection reagent (Thermofisher #13778150) to 50 μL Opti-MEM in another. The two tubes were then mixed and incubated at room temperature for 15 minutes. Cells were trypsinized, resuspended in 10% FCS DMEM, and the desired cell number added to each well. The transfection mix was then added in a dropwise fashion to these cells and incubated for 48 hours. At that time, transfected cells were trypsinized, counted, and replated for the desired assay overnight (including interferon pretreatment); cells were infected 72 hours after the initial siRNA transfection. For a 6-well, all values were doubled (600,000 cells; 25 pmol siRNA in 100 μL OptiMEM, 8 μL RNAiMAX in in 100 μL OptiMEM).

### Interferon Pretreatment

Recombinant human IFN-gamma was obtained from ProSpec (#CYT-206) and resuspended in sterile distilled water to a concentration of 2 x 10^7 units/mL (equivalent to 1 mg/mL or 58.82 μM). Recombinant universal type I interferon (human IFN-alpha hybrid protein) was obtained from PBL Assay Science (#11200) and resuspended in sterile distilled water to a concentration of 2 x 10^7 units/mL (equivalent to 0.128 mg/mL or 6.575 μM). A minimum of 4 hours after plating, cells were pretreated or mock pretreated with the desired amount of interferon alpha or gamma for 16 hours (flow cytometry, luciferase, plaque assay) or 24 hours (western blot, immunoprecipitation, microscopy). Interferon alpha or gamma treated cells in Figure 1 received the indicated molarity of interferon. All other interferon alpha or gamma treatments received 500 units (96-well) or 5000 units (24-well) interferon suspended in 30 μL serum-free DMEM. Mock treated cells received 30 μL serum-free DMEM.

### Flow Cytometry of Infected Cells

Cells in a 96-well plate were infected by adding 30 μL of serum-free DMEM containing the desired multiplicity of infection (MOI) of virus. At the desired timepoint (24h, 48h, or 72h), the media was removed, cells washed with PBS, and trypsinized. They were then resuspended in a final concentration of 2% paraformaldehyde in a 96-well V-bottom plate. After 15 minutes of fixation, cells were pelleted at 1000 x g. Cells expressing a fluorescent reporter were resuspended in 200 μL PBS containing 0.1% Pluronic F-68 (now known as Poloxamer 188, Thermofisher #24040032) before running on an Invitrogen Attune NxT Flow Cytometer. Cells requiring antibody staining were resuspended in 150 μL blocking buffer (PBS with 5% FCS and 0.05% sodium azide) for 5 minutes, then spun down and resuspended in primary antibody staining solution for 30 minutes (PBS with 5% FCS, 0.5% saponin, 0.05% sodium azide). The following antibodies were used, all 1:1000 dilution except where noted: HCoV-OC43 mouse anti-nucleocapsid (Sigma #MAB9013), HCoV-229E mouse anti-nucleocapsid (Sino Biological #40640-MM11), HCoV-NL63 rabbit anti-nucleocapsid (Sino Biological #40641-T62), influenza A virus mouse anti-nucleocapsid (BEI Resources #NR-19868, 1:5000), adenovirus mouse anti-hexon conjugated to FITC (Thermofisher #MA1-7329, 1:500). Cells requiring a secondary antibody for fluorescence (all except adenovirus) were spun down, supernatant removed, washed with staining solution once, and then resuspended in staining solution containing a 1:1000 dilution of secondary antibody for 30 minutes, either goat anti-mouse conjugated to AF488 (Thermofisher #A-11029) or goat anti-rabbit conjugated to AF488 (Thermofisher #A-11034). After washing, cells were resuspended in 200 μL PBS containing 0.1% Pluronic F-68 and run on an Invitrogen Attune NxT Flow Cytometer. Data was analyzed in FlowJo v10.10. An example of the gating strategy is below:

### Plaque Assay

100,000 cells (already transfected with siRNAs as above) were plated in a 24 well plate for 24 hours to make a confluent monolayer of cells including treatment with or without interferon gamma as needed for 16 hours. Coronavirus stocks were diluted in 10-fold serial dilutions in serum-free DMEM, and three dilutions selected based on the titer of the virus stock. Cells were washed with PBS, and 200 μL of virus inoculum used to infect each well at 34°C. Cells were rocked every 15 minutes for an hour, and the inoculum removed. 425 μL of an overlay containing 1% methylcellulose in 2% FCS DMEM was then added and incubated for 4 days. The overlay was then removed, and cells fixed in a final concentration of 2% paraformaldehyde in PBS for 15 minutes. After this, cells were washed with PBS. For viruses expressing an eGFP reporter (rHCoV-OC43/eGFP, rSARS-CoV-2/eGFP), plates were imaged at this step (see below). For viruses requiring staining, (HCoV-229E, HCoV-NL63), the monolayer was first incubated for 5 minutes in PBS containing 5% FCS and 0.05% sodium azide, washed again in PBS, then stained for 30 minutes in 5% FCS, 0.5% saponin, and 0.05% sodium azide containing a 1:1000 dilution of the following antibodies: HCoV-229E mouse anti-nucleocapsid (Sino Biological #40640-MM11), HCoV-NL63 rabbit anti-nucleocapsid (Sino Biological #40641-T62). Cells were washed in PBS, then stained for 30 minutes in 5% FCS, 0.5% saponin, and 0.05% sodium azide containing a 1:1000 dilution of the appropriate secondary antibody: goat anti-mouse conjugated to AF488 (Thermofisher #A-11029) or goat anti-rabbit conjugated to AF488 (Thermofisher #A-11034). Cells were then washed and resuspended in PBS before imaging. Plaque assays were imaged on a Thermofisher EVOS Imaging System in the 488 channel, using a 2X objective to take 20 individual images to encompass the full area of each well, and then stitched together to make a final image. Plaques were counted manually by eye from the final stitched image to determine titer.

Plaque size was quantified as follows. The final stitched image was first cropped to eliminate signal from the edge of the well. This was done by creating a circle in ImageJ to encompass the cells in the well but exclude the well edge and the rest of the image which did not contain cells, then deleting all signal outside of the circle. This image was then converted for use in Imaris (Oxford Instruments) using ImarisFileConverter 11.0.1 and opened in Imaris 10.1. A program was made to analyze the plaque size of each virus as follows, with separate programs made for each virus. The largest plaque was measured manually and rounded to the nearest 100 μm. Plaques were then defined as 2D surfaces though background subtraction (local contrast) with the “diameter of largest sphere” set to that number. Threshold was set manually, and additional filters added for circularity (to eliminate aberrant detection of irregular signal not originating from a plaque) and minimum voxel size (to eliminate signal from single cells or otherwise not originating from a multiple-cell plaque). The area of each plaque (surface) was then exported and graphed in Prism.

Since plaques for SARS-CoV-2 were larger and contained areas of dead cells in the middle of the plaque with no fluorescent signal, a separate method was required to determine the size of SARS-CoV-2 plaques accurately. The size of SARS-CoV-2 plaques was determined by training a machine learning algorithm in Imaris against manually marked plaques as signal, and the rest of the plate as background. This was done separately for “No IFN” and “IFN-gamma” conditions, due to the low signal of the SARS-CoV-2 eGFP reporter and the stark difference in plaque morphology between the two conditions. The area of each plaque (surface) was then exported and graphed in Prism.

### FLAG Immunoprecipitation

Cells expressing FLAG-RNF213 were pretreated with IFN-gamma for 24 hours, followed by infection with rHCoV-OC43/eGFP-NSP3 or rSARS-CoV-2/eGFP-NSP3 at MOI 3 for another 24 hours. Cells were harvested by trypsinization, resuspended in 10% FCS DMEM, washed in PBS, then lysed in 200 μL lysis buffer containing 50 mM Tris pH 8, 150 mM NaCl, 1% Triton, and EDTA-free protease inhibitor cocktail (Roche #11836170001) for 10 minutes on ice. Lysed cells were pelleted for 30 minutes at max speed at 4°C. 20 μL (10%) of the supernatant was reserved as input, the rest was used for the IP. Protein G magnetic beads (Thermofisher “Dynabeads” #10004D) were incubated with 4 μg (4 μL of 1 mg/ml) mouse anti-FLAG antibody (Clone M2, Sigma #F1804) at room temperature for 2 hours in lysis buffer. After washing in lysis buffer, beads were added to the rest of the cell supernatant and incubated overnight at 4°C on a rotator. The next day, cells were washed in lysis buffer 5 times using a magnet, then resuspended in 10 μL 1X NuPAGE LDS Sample Buffer (Thermofisher # NP0008) containing 2% beta-mercaptoethanol. After denaturing at 70°C for 10 min, the supernatant was isolated from the beads and western blot performed.

### Whole Cell Lysis

Cells were trypsinized, resuspended in 10% FCS DMEM, washed in PBS, and lysed in RIPA buffer (Sigma #R0278) and EDTA-free protease inhibitor cocktail (Roche #11836170001) for 10 minutes on ice. Lysed cells were pelleted for 30 minutes at max speed at 4°C, and supernatant added to 4X NuPAGE LDS Sample Buffer (Thermofisher # NP0008) to a final concentration of 1X and 2% beta-mercaptoethanol. Proteins were denatured at 70°C for 10 min, and western blot performed.

### Western Blot

Samples were run on a NuPAGE Bis-Tris Midi Protein gel, 4-12% acrylamide, 1.0 mm (Thermofisher #WG1403) at 150 constant volts until the dye front reached the bottom (about an hour) and transferred to a nitrocellulose membrane (Cytiva #10600002) for 30 minutes at 25 constant volts in a SureLock Tandem Midi Blot Module (Thermofisher #STM2001). Membranes were blocked in Intercept PBS Blocking Buffer (Licor # 927-70001) for 15 minutes and incubated overnight at 4°C with a primary antibody (see below) diluted in Intercept blocking buffer. After washing 3 times for 5 minutes in PBS with 0.1% Tween-20 (PBST), membranes were incubated with a secondary antibody diluted 1:5000 in blocking buffer for 1 hour (either Licor Goat anti-rabbit 800 #926-32211, Goat anti-mouse 800 #926-32210, Goat anti-rabbit 680 #926-68071, Goat anti-mouse 680 #926-68070, or Goat anti-mouse 549 #926-54010). After washing 3 times for 5 minutes in PBST, membranes were incubated again (as indicated) with an actin antibody pre-conjugated to a 488 fluorophore (Proteintech #CL488-60008) for 1 hour and washed 3 times for 5 minutes in PBST. Blots were then scanned on a Licor Odyssey M imager in the 800, 700, 520, and/or 488 channels. Primary antibodies and concentrations used include: mouse anti-RNF213 (Santa Cruz #sc-293391, 1:500), mouse anti-OC43 N (Sigma #MAB9013, 1:1000), rabbit anti-HSP90 (Proteintech #13171-1-AP, 1:5000), mouse anti-FLAG (Sigma #F1804, 1:2000), rabbit anti-GFP (Proteintech #50430-2-AP, 1:2000), mouse anti-actin pre-conjugated to 488 (Proteintech #CL488-60008, 1:5000).

### Metabolic Labeling and Isolation of Viruses

Cells were initially plated overnight in DMEM with 10% FCS in a 6 well (600,000 cells). The next day, cells were washed in PBS, and cells were infected at MOI 5 with HCoV-OC43 in 250 μL serum-free DMEM. At 1 hpi, DMEM with 10% FCS was added to a final volume of 2 ml. At 4.5 hpi, cells were washed with PBS and media changed to 1 mL serum-free DMEM without methionine or cysteine (Thermofisher #21013024). After starvation for 30 minutes, 1 mM L-Azidohomoalanine (AHA), a methionine analog containing an azido moiety, was added overnight. At 24 hours post infection, viral supernatants were harvested and clarified by centrifugation at 1000 x g for 10 minutes. Supernatants (1 mL) were then floated on top of 4 mL of 30% sucrose in a 5 mL ultraclear ultracentrifuge tube (Beckman Coulter #344057) and ultracentrifuged in an SW-55 Ti rotor (Beckman Coulter #342194) for 3 hours at 30,000 RPM. The supernatant was discarded and viral pellet lysed and resuspended in 50 μL of 50 mM Tris-HCl pH 8.0 and 1% sodium dodecyl sulfate (SDS). Virions labeled with AHA were fluorescently labeled with AZDye 680 Alkyne (Vector Laboratories #CCT-1514) using the “Click-&-Go® Protein Reaction Buffer Kit” (Vector Laboratories #CCT-1262) according to the manufacturer’s instructions, followed by methanol precipitation. Briefly, to the final 200 μL click reaction mixture was added 600 μL methanol, 150 μL chloroform, and 400 μL water. After centrifuging at maximum speed for 5 minutes, the aqueous top layer was removed. 500 μL methanol was further added to pellet the protein, and after centrifuging again the supernatant was discarded. This was repeated once more to clean the pellet. The pellet was air-dried for 10 minutes, then resuspended in 10 μL 1X NuPAGE LDS Sample Buffer (Thermofisher # NP0008) containing 2% beta-mercaptoethanol. After denaturing at 70°C for 10 min, the sample was run on a NuPAGE Bis-Tris Protein Gel, 4-12%, 1.0 mm (Thermofisher #NP0323BOX). The gel was imaged by direct scanning on a LiCor Odyssey M imager in the 680 channel. After scanning, the gel was transferred to a membrane and western blot performed to verify the identity of each band.

### Generation of Recombinant Viruses

Recombinant viruses were made based on a previously published system using a circular polymerase extension reaction (CPER)^8^. Briefly, the sequence of HCoV-OC43 was split into 8 different fragments and cloned into plasmids, as well as a linker plasmid containing a cytomegalovirus (CMV) promoter, bovine growth hormone (BGH) poly(A) signal, and hepatitis delta virus ribosome ribozyme. Each fragment contained at least 34 bp of overlap with the preceding and proceeding fragment. Each of these fragments plus linker were PCR amplified, and gel extracted. 0.1 pmol of each fragment were then joined and amplified using a CPER PCR using PrimeStar Gxl DNA polymerase (Takara Bio #R050A) in a 50 μL reaction and the following program: 98 °C for 1 minute; 20 cycles of 98 °C for 10 seconds, 55 °C for 15 seconds, and 68 °C for 25 minutes; final extension 68 °C for 25 minutes; hold at 4 degrees. Half of this reaction (25 μL) was then transfected into 200,000 293T cells in a 6 well plate using Lipofectamine 2000 (Thermofisher # 11668019) as indicated by the manufacturer. After cytopathic effect was observed (usually around 7 days post transfection), the supernatant was harvested and virus propagated.

Recombinant HCoV-OC43 viruses were produced by cloning the following modifications into the aforementioned plasmids.

NS2 Replacement in HCoV-OC43: In fragment 6, the first 13 amino acids (MAVAYADKPNHFI) in NS2 were retained, as well as the final 27 amino acids and stop codon (KSFHFRKACQNLDCNCLGFYESSVEEY*). The rest of the amino acid sequence was replaced with the following reporters:

rHCoV-OC43/eGFP: Replaced with eGFP
rHCoV-OC43/M-HiBiT: Replaced with the sequence of HCoV-OC43 membrane protein except for the first methionine (SSKTTP…RNNI), followed by a GGSG linker, followed by HiBiT (VSGWRLFKKIS).
rHCoV-OC43/NLuc-NSP3/eGFP: Replaced with eGFP
NSP3 fusions in HCoV-OC43: NSP3 fusions were performed as previously described for MHV^9^. In fragment 1, the NSP2 gene was removed, except for the first two amino acids (VK). eGFP or Nanoluc-Luciferase were then inserted, followed by another YRGVK sequence before the regular NSP3 sequence. For example: [NSP1]GYRG[NSP2]VK[eGFP]MVSK…DELYK[NSP2]YRGVK[NSP3]GRRV…

rHCoV-OC43/eGFP-NSP3: eGFP sequence in place of NSP2
rHCoV-OC43/NLuc-NSP3/eGFP: Nano-luciferase sequence in place of NSP2, as well as eGFP in place of NS2 in fragment 6 as in the previous section.

Plasmids containing SARS-CoV-2 (NC_045512) split into 10 fragments, plus a linker plasmid, were purchased and synthesized by Thermofisher, with similar overlap (at least 34 bp) generated between fragments. Fragment PCR and CPER were performed identically to that of HCoV-OC43. Recombinant viruses were produced by cloning into these plasmids:

- RF7 replacement in SARS-CoV-2: ORF7 was replaced with eGFP, retaining the first 13 amino acids (MKIILFLALITLA) and last 27 amino acids (ELYSPIFLIVAAIVFITLCFTLKRKTE).
- fusion in SARS-CoV-2: NSP2 was deleted and replaced with eGFP, with only the first two amino acids being retained to keep the LXGGXX cleavage site after NSP1, immediately continuing with the NSP3 sequence: [NSP1]ELNGG[NSP2]AY[eGFP]MVSK…DELYK[NSP3]APTKVTF.

### Immunofluorescence Microscopy

8-well chambered coverglass slides (Thermofisher #12-565-338, Nunc Lab-Tek II borosilicate glass 1.5) were treated with gelatin (Sigma #G1393) for 1 hour at 37 °C. After aspirating, 20,000 cells were plated overnight in 10% FCS DMEM. Cells were treated with interferon for 24 hours, followed by infection as needed at MOI 3. If needed, lipid droplets were labeled by incubation with Lipid-Deep Red (Dojindo #LD04-10) at a final concentration at 1 nmol / μL for 2 hours before fixation. Media was aspirated, cells washed in PBS, then fixed in a final concentration of 2% paraformaldehyde in PBS for 15 minutes. All subsequent solutions (except PBS) were filtered through a 0.22 μm filter before using (Sigma #SCGP00525). After washing 3 times in PBS, cells were permeabilized with 1% triton in PBS for 5 minutes at room temperature. Cells were then washed 3 times in PBS, and blocked with PBS containing 30% FCS, 1% bovine serum albumen (BSA), and 0.01% Tween-20 for 15 minutes. After washing once in PBS, cells were incubated with a 1:500 dilution of primary antibody in PBS containing 1% BSA and 0.01% Tween-20 overnight at 4 °C. The next day, cells were washed 3 times in PBS and incubated with a 1:1000 dilution of secondary antibody and 2.5 μg/mL Hoechst 33342 stain (Abcam #ab145596) in PBS containing 1% BSA and 0.01% Tween-20 for 1 hour at room temperature. After washing 3 times in PBS, cells were left in PBS and imaged using a DeltaVision OMX SR imaging system and a 60X widefield oil immersion objective (Olympus), with exposure times of 100 ms in each channel, and laser power between 0.2% and 5% depending on the brightness of the target to prevent saturation of the detector. After setting within an experiment, laser power was kept constant between all samples. Images were aligned and deconvolved using softWoRx (DV files) and later cropped to show points of interest within the Fiji package for ImageJ2 (v2.16/1.54p). Cropped files were saved separately from the original data as TIFF files. Co-localization analysis was conducted using the Fiji package for ImageJ2 (v2.16/1.54p), and the BioImaging And Optics Platform (BIOP) JACoP plugin.

The following primary antibodies were used for microscopy: mouse anti-RNF213 (Sigma #MABN2510), mouse anti-FLAG (Sigma #F1804), rabbit anti-calnexin (Proteintech #10427-2-AP), rabbit anti-GM130 (Proteintech #11308-1-AP), sheep anti-TGN46 (Biorad #AHP500G), rabbit anti-MAVS (Proteintech 14341-1-AP), LC3A/B (Cell Signaling #12741T), EEA1 (Cell Signaling #3288S), CD63 (Abcam #ab252919), Clathrin (Cell Signaling #4796S), Caveolin (Santa Cruz #sc-53564), LAMP1 (Cell Signaling #9091S). All primary antibodies were used with secondaries raised in goat, except for experiments involving the sheep anti-TGN46 antibody, which used exclusively secondaries raised in donkey. The following secondary antibodies were used: Goat anti-mouse 488 (Thermofisher #A11029), Goat anti-rabbit 488 (Thermofisher #A11034), Goat anti-mouse 568 (Thermofisher #A11004), Goat anti-rabbit 568 (Thermofisher #A11036), Goat anti-mouse 647 (Thermofisher #A21235), Goat anti-rabbit 647 (Thermofisher #A21245), Donkey anti-sheep 647 (Thermofisher #A21448), Donkey anti-mouse 568 (Thermofisher #A10037).

### Luciferase Assays

#### Nano-luciferase (lysed)

Following infection with VSV expressing a Nano-luciferase reporter (Fig 1), reporter expression was quantified by lysing cells and determining luciferase activity. Cells in a 96 well black plate were washed with PBS and lysed in 30 μL 1X lysis buffer (diluted from 5X, Promega #E1531) for 15 minutes. Promega Nano-Glo Luciferase reagent was diluted 1:50 in its buffer according to the manufacturer’s instructions (Promega Nano-Glo Luciferase Assay System #N1150). 30 μL diluted luciferase reagent was added to the lysed cells and incubated at room temperature for 5 minutes, then activity read on a Promega GloMax Navigator luminometer plate reader.

#### Complete Nano-luciferase (live)

Following infection with HCoV-OC43 NLuc-NSP3 (Fig 6F), reporter expression was quantified over time in live cells (without lysis). Cells in a 96 well black plate were washed with PBS, and infected with rHCoV-OC43/NLuc-NSP3/eGFP at MOI 1 in 10% FCS DMEM containing a 1:100 dilution of the Nano-Glo Endurazine Substrate according to the manufacturer’s instructions (Promega #N2571). Controls added concurrently with infection included 200 μM Molnupiravir (EIDD-2801, Sigma #SML2873), or 500 μg/mL cycloheximide (Sigma #C4859). Infection proceeded at 34 °C and was read hourly on a Promega GloMax Navigator luminometer plate reader.

#### HiBiT/LgBiT Reconstitution (live)

Cells were transduced with a lentivirus expressing LgBiT from a CMV promoter. Cells were then infected in a 96 well plate with rHCoV-OC43/M-HiBiT at MOI 3 in 10% FCS DMEM containing a 1:100 dilution of the Nano-Glo Endurazine Substrate according to the manufacturer’s instructions (Promega #N2571). Controls added concurrently with infection included 200 μM Molnupiravir (EIDD-2801, Sigma #SML2873), 500 μg/mL cycloheximide (Sigma #C4859), or 25 mg/mL soluble heparin (Sigma #H3393). Infection proceeded at 34°C. HiBiT/LgBiT reconstitution of luciferase activity was read hourly on a Promega GloMax Navigator luminometer plate reader.

### qPCR of Viral RNA

A549 cells were seeded in a 24 well plate (100,000 cells), pretreated with interferon-gamma for 16 hours as above, and infected with HCoV-OC43 at MOI 1. Normal human serum (15%) was added at 4 hpi to block spread (Sigma #S1-M). At 12 hpi, cells were washed in PBS and lysed in 1 mL Trizol (Thermofisher #15596018). RNA was extracted according to the manufacturer’s instructions. Briefly, 200 μL of chloroform (Sigma #C2432) was added, solution vortexed, and allowed to separate for 1 minute. This was then centrifuged at max speed for 15 minutes at 4°C. The supernatant was removed, 1 μL glycogen added (Roche #10901393001), and an equal volume (∼300 μL) of isopropanol added (Sigma I9516). After incubating at room temperature for 10 minutes, this was centrifuged at max speed for 10 minutes at 4°C. The supernatant was removed, and pelleted RNA washed with 500 μL of 70% ethanol (made fresh from 100% ethanol from Decon Labs #2716 and sterile/RNase/DNase free water from Thermofisher #46-000-CI). This was centrifuged at max speed for 5 minutes at 4°C, supernatant removed, pellet dried, and resuspended in 20 μL water (Thermofisher #46-000-CI).

A reverse transcription reaction was performed using 1 μg of RNA and the SuperScript III First-Strand Synthesis System (Thermofisher #18080051) using the provided random hexamers according to the manufacturer’s protocol. qPCR was performed using a TaqMan Fast Advanced Master Mix (Thermofisher #4444557) according to the manufacturer’s protocol using the following combination of primers and probes. We used a duplexed system in which either the OC43 genomic RNA or subgenomic nucleocapsid mRNA could be detected in one channel (FAM dye), and actin in another channel (JOE dye). We purchased the following taqman probes from IDT: Actin (/56-JOEN/ATCAAGATC/ZEN/ATTGCTCCTCCTGAGCGC/3IABkFQ/); OC43 N (/56-FAM/CAGTAGTAG/ZEN/AGCGTCCTCTGGAAATCGTTC/3IABkFQ/). For actin amplification, we used the following primers: Actin Forward (CCTGGCACCCAGCACAAT), Actin Reverse (GCCGATCCACACGGAGTACT). For OC43 genomic and subgenomic RNA, we used a conserved reverse primer in the OC43 N gene (TTGAGCTCTTCTACCCCTG) with the aforementioned probe. To detect specifically the subgenomic RNA, we used a sequence in the OC43 leader, which is present in subgenomic RNAs, but not in genomic RNAs (CGCTTCACTGATCTCTTGTTAG). To detect genomic RNA, we used a sequence in the membrane protein which precedes nucleocapsid in the genome, but not subgenomic RNAs (CATCAACCCAAAAGGGTTCTG). qPCR was run in 40 cycles on Applied Biosystems StepOnePlus System and thresholded in Quantstudio to determine CT values. All three biological replicates were run simultaneously on the same plate. Data was analyzed using the ΔΔCT method in Excel. Briefly, one empty vector control in one replicate was set to a ΔCT value of 0. The difference in CT value for all other samples was calculated (including the EV controls from that replicate, and all EV controls from other replicates). The ΔCT for actin was then subtracted from the ΔCT for OC43 mRNA or genome, producing a ΔΔCT value of OC43 mRNA to actin, or OC43 genome to actin. The calculation 2^(ΔΔCT) was then made for all values such that one control was set to 1, and the relative value of each other sample and replicate was calculated.

### Data Analysis and Graphs

Primary data (e.g. FlowJo, ImageJ, Promega GloMax Luminometer) was exported to a Microscoft Excel file. This data was copied into Prism v10.6.1 to generate graphs. Statistical tests were performed in Prism. In experiments with only two conditions (e.g. EV vs RNF213 KO) an unpaired, two-tailed, t-test using a normal (Gaussian) distribution assumption was performed. For experiments with conditions varying interferon treatment (No IFN vs IFN-gamma) and RNF213 (si-NT vs si-RNF213) an “ordinary” 2-way ANOVA was performed unless otherwise indicated in the figure legend. In both cases, the resulting p-values are displayed on each graph. For experiments using CRISPR-edited SCCs, data is displayed on each graph and statistics calculated with each single-cell clone of a cell line treated as an individual biological replicate (one representative experiment shown, +/- standard deviation). For experiments using siRNA transfection, data is displayed on each graph and statistics calculated as the average of 3 biological replicates (+/- standard deviation).

### Schematics

The schematic in Figure 1D was generated with the assistance of Gemini (Google) and was reviewed and edited by the authors for scientific accuracy. Other schematics were made in Microsoft Powerpoint.

### Example of Gating Strategy for Flow Cytometry

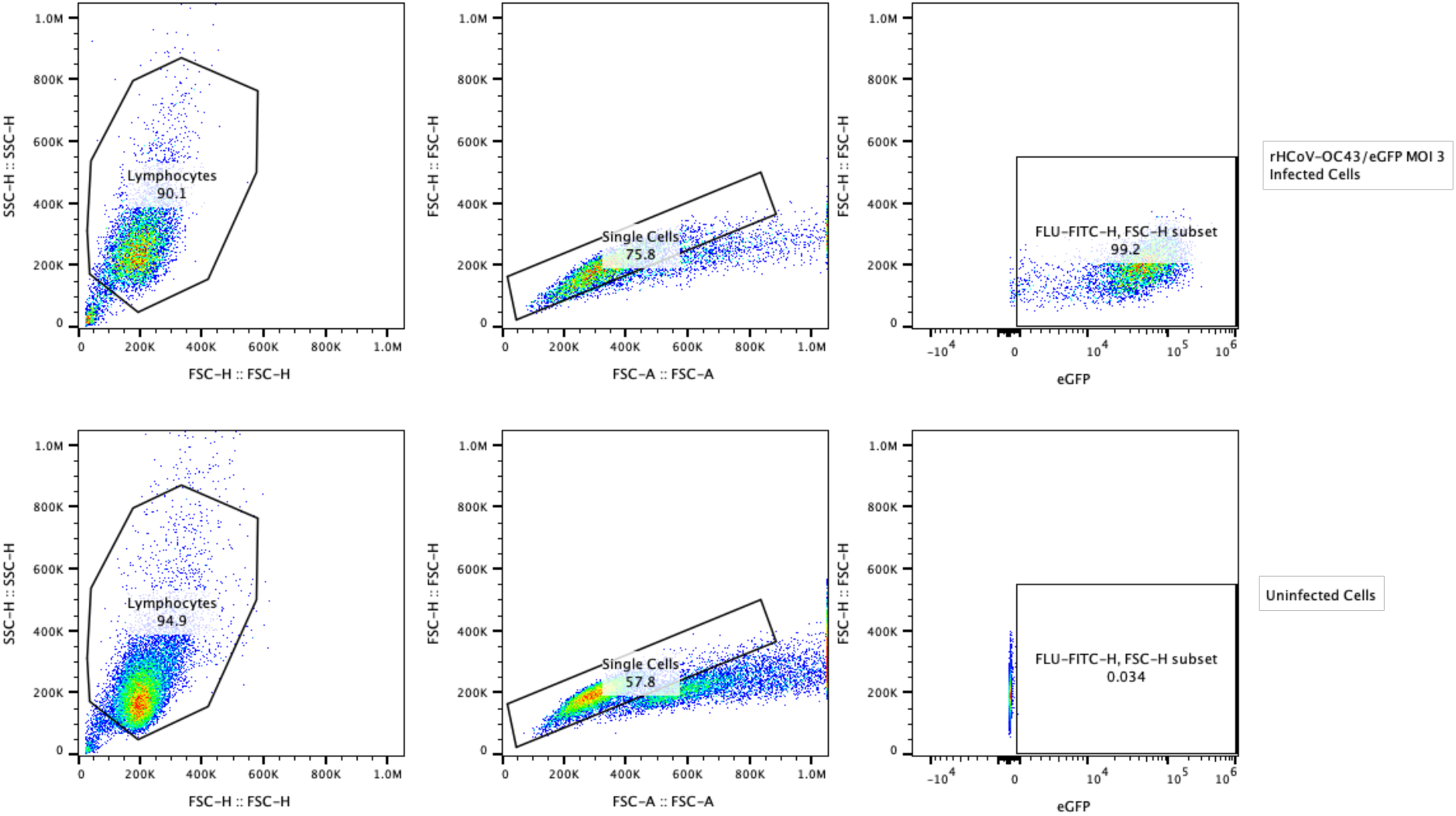

### Table of All Antibodies Including Lot Numbers

| Host | Target | Fluorophore | Supplier Name | Catalog Number | Clone Name | Lot Number |
| --- | --- | --- | --- | --- | --- | --- |
| Mouse anti- | HCoV-229E Nucleocapsid |  | Sino Biological | 40640-MM11 | Not given | HB150C2803-B |
| Rabbit anti- | HCoV-NL63 Nucleocapsid |  | Sino Biological | 40641-T62 | Polyclonal | HD15DE3011 |
| Mouse anti- | HCoV-OC43 Nucleocapsid |  | Millipore Sigma | MAB9013 | 542-7D | 3754724 |
| Mouse anti- | FLAG |  | Millipore Sigma | F1804 | M2 | 375542 |
| Mouse anti- | Actin | 488 | Proteintech | CL488-60008 | 7D2C10 | 21023753 |
| Mouse anti- | RNF213 |  | Santa Cruz | sc-293391 | 5C12 | F0524 |
| Mouse anti- | RNF213 |  | Millipore Sigma | MABN2510 | 1C9 | 4244929 |
| Rabbit anti- | HSP90 |  | Proteintech | 13171-1-AP | Polyclonal | 126494 |
| Rabbit anti- | GFP |  | Proteintech | 50430-2-AP | Polyclonal | 176935 |
| Rabbit anti- | Calnexin |  | Proteintech | 10427-2-AP | Polyclonal | 106234 |
| Rabbit anti- | GM130 |  | Proteintech | 11308-1-AP | Polyclonal | 120702 |
| Sheep anti- | TGN46 |  | Biorad | AHP500G | Polyclonal | 164005 |
| Rabbit anti- | MAVS |  | Proteintech | 14341-1-AP | Polyclonal | 76926 |
| Rabbit anti- | LC3A/B |  | Cell Signaling | 12741T | D3U4C | 5 |
| Rabbit anti- | EEA1 |  | Cell Signaling | 3288S | 3288S | 10 |
| Rabbit anti- | CD63 |  | Abcam | ab252919 | EPR22458-280 | GR3270794-4 |
| Rabbit anti- | Clathrin |  | Cell Signaling | 4796S | D3C6 | 6 |
| Mouse anti- | Caveolin |  | Santa Cruz | sc-53564 | 7C8 | K1922 |
| Rabbit anti- | LAMP1 |  | Cell Signaling | 9091S | D2D11 | 6 |
| Mouse anti- | Adenovirus Hexon | 488 | Thermofisher | MA1-7329 | D36J | YB3833843 |
| Mouse anti- | Influenza NP A/CA/04/09 H1N1 |  | BEI Resources | NR-19868 | 2F4 | 70027831 |
| Goat anti- | Mouse | 488 | Thermofisher | A-11029 |  | 3268353 |
| Goat anti- | Rabbit | 488 | Thermofisher | A-11034 |  | 3124362 |
| Goat anti- | Mouse | 568 | Thermofisher | A11004 |  | 2633533 |
| Goat anti- | Rabbit | 568 | Thermofisher | A11036 |  | 2720366 |
| Goat anti- | Mouse | 647 | Thermofisher | A21235 |  | 2482945 |
| Goat anti- | Rabbit | 647 | Thermofisher | A21245 |  | 2641997 |
| Donkey anti- | Mouse | 568 | Thermofisher | A10037 |  | 2674370 |
| Donkey anti- | Sheep | 647 | Thermofisher | A21448 |  | 2563882 |
| Goat anti- | Mouse | 549 | Licor | 926-54010 |  | D50206-01 |
| Goat anti- | Rabbit | 680 | Licor | 926-68071 |  | D50729-04 |
| Goat anti- | Mouse | 680 | Licor | 926-68070 |  | D21115-35 |
| Goat anti- | Rabbit | 800 | Licor | 926-32211 |  | D50805-03 |
| Goat anti- | Mouse | 800 | Licor | 926-32210 |  | D50826-03 |

